# Targeting SERCA2 in organotypic epidermis reveals MEK inhibition as a therapeutic strategy for Darier disease

**DOI:** 10.1101/2023.03.07.531620

**Authors:** Shivam A. Zaver, Mrinal K. Sarkar, Shaun Egolf, Jonathan Zou, Afua Tiwaa, Brian C. Capell, Johann E. Gudjonsson, Cory L. Simpson

## Abstract

Mutation of the *ATP2A2* gene encoding sarco-endoplasmic reticulum calcium ATPase 2 (SERCA2) was linked to Darier disease more than two decades ago; however, there remain no targeted therapies for this disorder causing recurrent skin blistering and infections. Since *Atp2a2* knockout mice do not phenocopy its pathology, we established a human tissue model of Darier disease to elucidate its pathogenesis and identify potential therapies. Leveraging CRISPR/Cas9, we generated human keratinocytes lacking SERCA2, which replicated features of Darier disease, including weakened intercellular adhesion and defective differentiation in organotypic epidermis. To identify pathogenic drivers downstream of SERCA2 depletion, we performed RNA sequencing and proteomic analysis. SERCA2-deficient keratinocytes lacked desmosomal and cytoskeletal proteins required for epidermal integrity and exhibited excess MAP kinase signaling, which modulates keratinocyte adhesion and differentiation. Immunostaining patient biopsies substantiated these findings with lesions showing keratin deficiency, cadherin mis-localization, and ERK hyper-phosphorylation. Dampening ERK activity with MEK inhibitors rescued adhesive protein expression and restored keratinocyte sheet integrity despite SERCA2 depletion or chemical inhibition. In sum, coupling multi-omic analysis with human organotypic epidermis as a pre-clinical model, we found that SERCA2 haploinsufficiency disrupts critical adhesive components in keratinocytes via ERK signaling and identified MEK inhibition as a treatment strategy for Darier disease.

## INTRODUCTION

Though advances in gene sequencing have facilitated the diagnosis of rare inherited disorders and enhanced our understanding of their pathogenic mechanisms (1–3), most genetic skin diseases lack proven, rational molecular therapies (4, 5). Mutation of the *ATP2A2* gene, which encodes sarco-endoplasmic reticulum calcium ATPase 2 (SERCA2), causes Darier disease (DD) (6, 7), a dermatologic disorder characterized by aberrant epidermal differentiation and impaired keratinocyte adhesion via desmosomes (8, 9), which manifests as recurrent skin blisters, erosions, and infections (10–13). Despite knowing the genetic etiology of this disease for more than twenty years, there are no FDA-approved therapies for DD and no drugs that directly target the pathogenic effects of SERCA2 deficiency have advanced to prospective clinical trials (14).

While systemic retinoids have been used off-label for treatment of DD (15, 16), their long-term toxicity and highly teratogenic nature limit utilization in patients suffering from this lifelong disorder (17, 18). Similarly, topical corticosteroids and antimicrobials can treat secondary inflammation and infection in localized lesions (14), but are not specifically aimed at the underlying disease pathophysiology and exhibit limited efficacy, especially in widespread disease that can lead to serious illness (12, 13, 19). Unfortunately, targeted ablation of the *Atp2a2* gene in mice did not replicate DD pathology, but instead led to increased age-related keratinocyte carcinomas (20, 21), a risk not borne out in patients.

We leveraged cultured human keratinocytes and organotypic epidermis (22) to establish a pre-clinical model of DD to better elucidate its pathogenesis and identify new potential therapeutic avenues to help direct future clinical trials. Using CRISPR/Cas9 gene editing (23, 24), we generated human keratinocytes having the first exon of *ATP2A2* disrupted and performed multi-omic analysis, which revealed inhibition of the mitogen-activated protein kinase (MAPK) pathway as a novel treatment strategy for DD.

## RESULTS

### Loss of SERCA2 in human keratinocytes impairs cytosolic calcium handling and reduces intercellular adhesive strength

Darier disease, an autosomal dominant disorder, has been linked to various *ATP2A2* gene mutations (6, 11, 25), most of which are predicted to cause haploinsufficiency of SERCA2, a calcium channel localized to the endoplasmic reticulum (ER) (26). SERCA2 function is essential for calcium-dependent assembly of intercellular junctions, particularly desmosomes, which maintain epidermal integrity (8, 9). As well, SERCA2 controls levels of cytosolic calcium, a master regulator of the terminal phase of keratinocyte differentiation (27, 28), termed cornification, in which cells in the upper epidermis degrade their nuclei and organelles to form the flattened outermost layers of the cutaneous barrier (29, 30). In DD patients, the loss of intercellular adhesion and dysfunctional keratinocyte differentiation manifest as blistering, thick scaling, and fissuring of the skin (10, 26).

Since *Atp2a2* KO mice did not phenocopy DD (20), pre-clinical studies of its pathogenesis have been limited and no therapies have advanced to clinical trials to achieve FDA approval (14). Given the cutaneous pathology of DD is limited to the epidermis, we used CRISPR/Cas9 to ablate the *ATP2A2* gene in keratinocytes (24), which can be grown into a fully stratified epidermis within an organotypic model (22). We targeted the first exon of *ATP2A2* (or a pseudogene as a control) in hTERT-immortalized human epidermal keratinocytes (THEKs) from the parental line N/TERT-2G (31); this allowed isolation of stable homozygous knockout (KO) and heterozygous (HET) cell lines harboring frameshift mutations that depleted SERCA2 as confirmed by immunoblotting cell lysates and immunostaining fixed cells (Fig. 1A-B).

**Figure 1.**
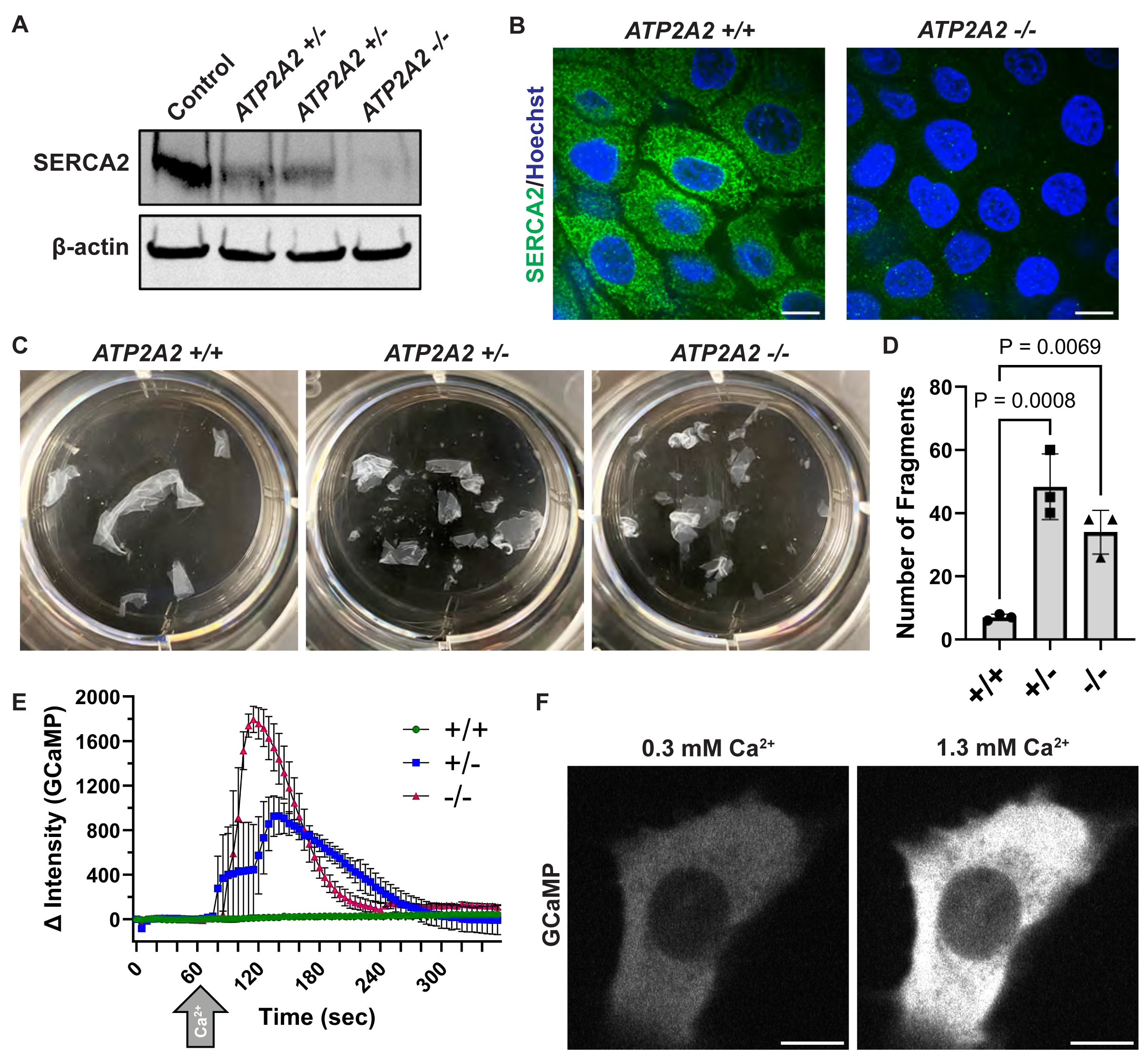
Loss of SERCA2 in human keratinocytes impairs cytosolic calcium handling and reduces intercellular adhesive strength. **(A)** Immunoblot of SERCA2 in lysates from *ATP2A2* wildtype (WT, +/+), heterozygous (HET, +/-), and homozygous knockout (KO, -/-) cells. hTERT-immortalized human epidermal keratinocytes (THEKs) were differentiated in E-medium for 72 h before harvesting lysates; data are representative of 3 independent experiments; β-actin is a loading control. **(B)** Immunofluorescence of SERCA2 (green) in WT and KO THEKs; images are representative of 14 independent high-powered fields (hpf) per genotype; Hoechst (blue) stains nuclei; scale bar = 10 μm. **(C)** Mechanical dissociation assay of monolayers from control (+/+), HET (+/-), and KO (-/-) cells grown in 1.3 mM CaCl_2_ for 72 h prior to using dispase to release intact monolayers; representative images of fragmented monolayers transferred into 6-well cell culture plates after mechanical stress are shown. **(D)** Bar graphs display the mean ± SD of the number of epithelial fragments with individual data points plotted for N=3 biological replicates; P-values are from one-way ANOVA with the Dunnett adjustment for multiple comparisons to control cells. **(E)** Change (Δ) in intensity of GCaMP in control (+/+), HET (+/-), and KO (-/-) THEKs from baseline at t = 0 s with addition of 1 mM CaCl_2_ at t = 60s (arrow); data are plotted as mean ± SEM from N=3 independent experiments per genotype. **(F)** Representative fluorescence images of GCaMP in HET cells in low CaCl_2_ (left) or high CaCl_2_ (right); scale bar = 10 µm.

Given the essential role of SERCA2 in assembling cell-cell junctions (8, 9), we assessed the mechanical integrity of control and SERCA2-deficient THEKs using an established dispase-based assay to quantify intercellular adhesive strength of epithelial cell monolayers (32, 33). We found that both HET and KO THEKs exhibit reduced integrity, reflected by increased fragmentation of epithelial sheets upon mechanical stress (Fig. 1C-D). We also used GCaMP, a ratiometric green fluorescent protein (GFP)-based genetically encoded calcium indicator (34, 35), to confirm that depletion of the ER-localized calcium ATPase altered cytosolic calcium handling. THEKs were transduced with GCaMP, then imaged by live microscopy during exposure to increased extracellular calcium. While control cells demonstrated minimal change in GCaMP intensity after treatment with calcium, KO cells experienced large increases in cytosolic calcium; HET cells exhibited an intermediate phenotype, as would be expected with haploinsufficiency of SERCA2-driven transport of calcium into the ER (Fig. 1D-E).

### Keratinocytes lacking SERCA2 exhibit transcriptomic alterations in mediators of epidermal adhesion and differentiation

To identify potential pathogenic drivers of DD in an unbiased manner, we assessed downstream consequences of SERCA2 depletion on the transcriptome in HET and KO keratinocytes. Bulk RNA sequencing revealed that HET cells exhibited significant changes in mRNA transcript profiles (Fig. 2A) with gene ontology (GO) analysis identifying alterations in gene subsets linked to intercellular adhesion, calcium regulation, and epidermal differentiation (Fig. 2B), consistent with known pathologic features of DD. However, we also identified a gene signature indicating elevation in epidermal growth factor receptor (EGFR) signaling and the downstream mitogen-activated protein kinase (MAPK) pathway, which are known to regulate epidermal differentiation and adhesion (36–40). We verified these SERCA2 deficiency-related changes in gene expression with quantitative reverse transcription PCR (RT-PCR) (Fig. 2C).

**Figure 2.**
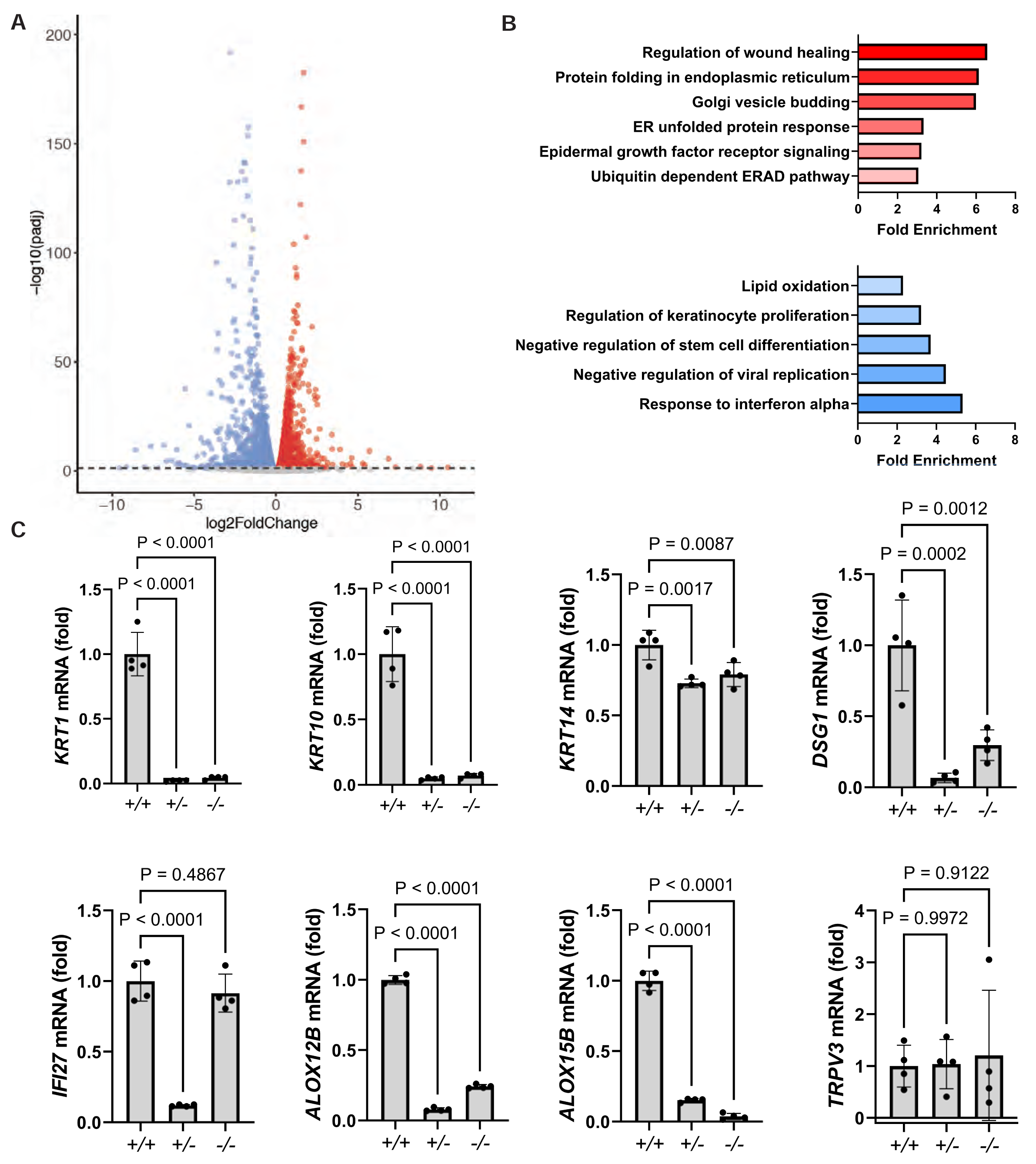
Keratinocytes lacking SERCA2 exhibit transcriptomic alterations in mediators of epidermal adhesion and differentiation. **(A)** Results from bulk RNA sequencing of SERCA2 HET versus control cells. A volcano plot depicts log_2_ fold-changes of genes significantly down-(blue) and up-regulated (red) with a cutoff (dashed line) of 0.05 for the adjusted P-value. **(B)** Gene ontology (GO) analysis of transcripts significantly altered (adjusted P-value≥0.05) in HET cells revealed up-regulation (red) in genes controlling ER stress and growth factor signaling and down-regulation (blue) of genes controlling epidermal differentiation and anti-viral response. **(C)** Control (+/+), HET (+/-), or KO (-/-) THEKs were differentiated in E-medium for 24 h, then mRNA transcripts were quantified by reverse transcription PCR (RT-qPCR). Bar graphs display the mean ± SD with plotted individual values from N=4 biological replicates; P-values are from one-way ANOVA with Dunnett adjustment for multiple comparisons to control cells.

Relative to controls, HET keratinocytes exhibited marked reduction in mRNA encoding keratin 1 (KRT1) and keratin 10 (KRT10) intermediate filament cytoskeletal proteins and the desmosomal cadherin desmoglein 1 (DSG1), all of which are known to exhibit differentiation-dependent expression in the suprabasal epidermal layers (41). In contrast, the effect of SERCA2 deficiency on mRNA encoding the primary keratin of the basal epidermal layer, KRT14, was much smaller. While GO analysis identified changes in some differentiation-associated calcium-binding proteins, the mRNA encoding TRPV3, a plasma membrane calcium channel recently linked to calcium influx during the final stage of keratinocyte differentiation (30), was unchanged in HET or KO cells. Interestingly, some genes exhibited disparate effects in KO versus HET lines, such as the interferon response gene *IFI27*, which was only depleted in HET cells.

The expression of lipoxygenase genes that drive membrane lipid modifications essential for terminal keratinocyte differentiation and epidermal barrier formation (42, 43) were also significantly reduced in SERCA2-deficient cells. Loss of ALOX12B, in particular, has been associated with autosomal recessive congenital ichthyosis (ARCI) (44), a group of inherited dermatologic diseases characterized by skin scaling and epidermal barrier defects due to defective keratinocyte maturation (43, 45), which shares clinical features with DD and also responds to retinoid therapy (17).

### Proteomic analysis of SERCA2-deficient keratinocytes reveals alterations in key regulators of tissue integrity and skin barrier formation

To determine if transcriptional changes in SERCA2-deficient keratinocytes translated to effects on protein expression, we performed label-free mass spectrometry (MS)-based identification of the proteome in two independent HET cell lines (Fig. 3A). This technique confirmed deficiency of SERCA2, itself, while the levels of other ER-localized proteins (SEC61B, SEC61A1) and an often-used housekeeping gene (GAPDH) were not significantly changed in HET cells compared to controls.

**Figure 3.**
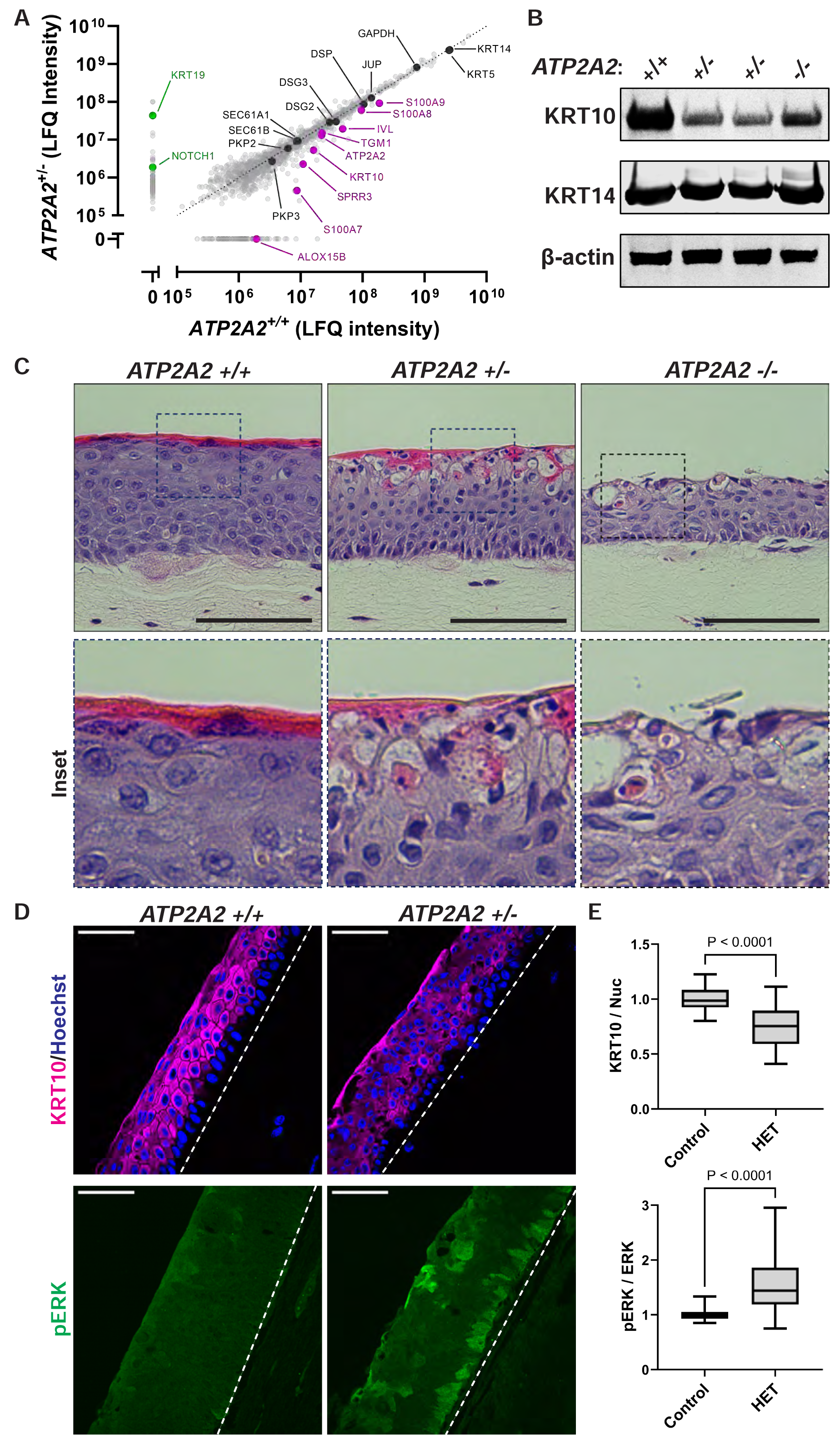
Proteomic analysis of SERCA2-deficient keratinocytes reveals alterations in key regulators of tissue integrity and skin barrier formation. **(A)** Label-free quantitative (LFQ) mass spectrometric-based comparison of the proteomes of control versus HET THEKs differentiated in E-medium for 72 h. Proteins highlighted in purple were absent or detected in significantly lower abundance in HET cells; proteins highlighted in green were detected in higher abundance in control THEKs; proteins highlighted in black were detected in similar abundance in control and HET cells. **(B)** Immunoblotting of KRT10 and KRT14 in lysates from control, HET, and KO THEKs differentiated in E-medium for 72 h; data are representative of 5 independent experiments; β-actin is a loading control. **(C)** H&E-stained tissue cross-sections from organotypic cultures grown from control, HET, or KO THEKs. HET and KO cultures display aberrant differentiation with the upper granular layers exhibiting vacuolization and loss of cohesion along with dyskeratotic cells having deeply pink cytoplasm and condensed nuclei indicative of aberrant cornification (insets); scale bar = 100 µm. **(D)** Immunostaining of KRT10 and pERK in tissue cross-sections from organotypic cultures of control (*ATP2A2*+/+) or HET (*ATP2A2*+/-) THEKs; scale bar = 50 µm; dashed line marks bottom of epidermis. **(E)** Quantification of tissue immunostaining of KRT10 (relative to Hoechst) and pERK (relative to total ERK); data are shown as a box plot of the 25^th^-75^th^ percentile with a line at the median from N≥25 non-overlapping hpf per condition from 4 experiments using 2 independent HET lines; control mean normalized to 1; P-values are from two-tailed unpaired Student’s t-test.

Proteomic analysis again revealed significant reduction in proteins that drive membrane lipid modifications (ALOX15B) and several members of the S100 family of calcium-binding proteins. As well, HET cells had a deficiency of involucrin (IVL) and transglutaminase 1 (TGM1), two key regulators of the cornified cell envelope, a highly cross-linked protein meshwork assembled at the inner surface of differentiated keratinocytes that makes the outermost epidermal layers water-impermeable (29, 43). The *TGM1* gene, in particular, has been linked to defective keratinocyte differentiation in patients with ARCI (46).

While many desmosome-associated proteins, including desmogleins (DSG2, DSG3), plakoglobin (JUP), desmoplakin (DSP), and plakophilins (PKP2, PKP3), exhibited little change in in SERCA2-deficient cells, their primary associated intermediate filament cytoskeletal component in the suprabasal epidermal layers, KRT10, was significantly reduced. If desmosomal complexes are not coupled to the keratin cytoskeleton, this dramatically compromises their ability to support intercellular adhesive strength (33). This finding was confirmed by immunoblotting HET and KO cell lysates, which exhibited reduced KRT10, while KRT14 was not notably altered (Fig. 3B).

Informed by our proteomic data, we aimed to determine if these alterations in SERCA2-deficient keratinocytes translated into pathogenic effects within a human tissue model. We grew KO and HET cells as organotypic skin cultures, which replicate the three-dimensional architecture of the stratified epidermis (22). Histologic analysis of epidermal cultures of HET cells revealed widening of intercellular spaces and marked disorganization of the upper keratinocyte layers undergoing the final stage of differentiation (Fig. 3C). Cornifying cells exhibited cytoplasmic vacuolization, impaired flattening, and retention of nuclei in the outermost keratinized layers (Fig. 3C, insets). These histologic defects occurred to a more extreme degree in KO cultures.

Consistent with our proteomic data, immunostaining SERCA2-deficient organotypic epidermis had reduced KRT10 in the suprabasal layers, which exhibited patchy expression of the cytoskeletal protein compared to the more uniform pattern seen in control cultures (Fig. 3D-E, upper panels). Since KRT10 is a major regulator of epidermal integrity and differentiation (47–49), a concomitant reduction of KRT10 in SERCA2-depleted epidermis could be responsible for the impaired intercellular adhesion and abnormal cornification seen in HET organotypic epidermis, which overlap with the pathologic features seen in DD biopsies.

We and others have previously shown that KRT10 expression is disrupted upon epidermal hyperactivation of MEK and ERK in the MAPK pathway (37, 50, 51). In fact, rises in cytoplasmic calcium, known to drive keratinocyte differentiation (27, 52), have been shown to trigger mitogen-independent MAPK signaling, which can be augmented by SERCA2 blockade (53). Further connecting this signaling pathway to DD, a DD-like disorder called Grover disease, characterized by similar defects in epidermal adhesion and keratinocyte cornification, has been linked to aberrant activation of MEK in patients (54, 55). These observations coupled with our transcriptomic and proteomic analyses led us to hypothesize that SERCA2 deficiency leads to aberrant activation of the MAPK pathway, which could represent a druggable target for treating DD (56).

Thus, we assessed whether hyper-activation of MEK, which functions in the MAPK pathway by activating extracellular-signal regulated kinase (ERK) via phosphorylation, could explain the defects in adhesion and differentiation seen in our DD tissue model. Reflecting increased MEK activity in SERCA2-deficient organotypic cultures, we found by immunostaining that phosphorylated ERK (pERK) was significantly elevated in HET tissues compared to control tissues (Fig. 3D-E, lower panels). These findings led us to examine whether suppressing MEK over-activation could rescue the phenotype of SERCA2 loss of function.

### Chemical inhibition of SERCA2 in keratinocytes and organotypic epidermis replicates features of DD pathology and induces ERK activation

To complement our results in SERCA2-deficient immortalized keratinocytes, we assessed the effect of a chemical inhibitor of SERCA2, thapsigargin (TG), on normal human epidermal keratinocytes (NHEKs). Treatment of NHEK monolayers with TG caused a significant increase in epithelial sheet fragmentation (Fig. 4A-B), which confirms the role of SERCA2 in promoting intercellular adhesion in primary human keratinocytes. Building on these findings, we treated mature organotypic epidermis grown from NHEKs with TG, which disrupted keratinocyte cohesion most notably between basal and suprabasal cell layers (Fig. 4C, inset), a pathologic feature of DD (55). Chemical inhibition of SERCA2 also disrupted keratinocyte differentiation causing retention of nuclei in the cornified layers (Fig. 4D, inset), a feature consistent with aberrant epidermal maturation (termed dyskeratosis) seen in DD (55, 57).

**Figure 4.**
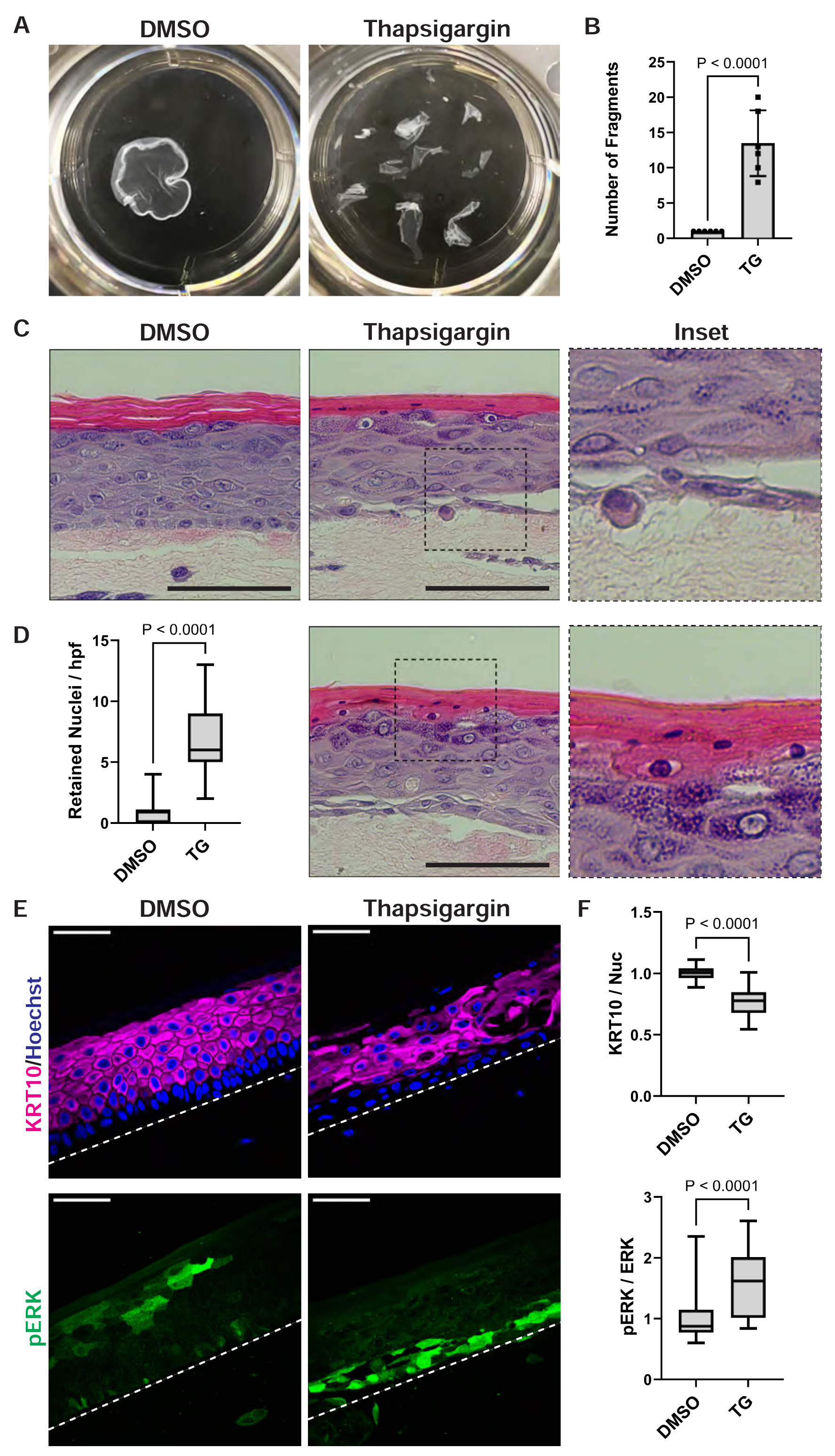
Chemical inhibition of SERCA2 in keratinocytes and organotypic epidermis replicates features of DD pathology and induces ERK activation. **(A)** Mechanical dissociation assay of confluent monolayers from THEKs cultured in 1.3 mM CaCl_2_ with DMSO or 1 μM thapsigargin (TG) for 24 h; representative images of fragmented monolayers transferred into 6-well cell culture plates are shown. **(B)** Bar graph displays the mean ± SD of the number of fragments with individual data points plotted for N=6 biological replicates; P-value is from two-tailed unpaired Student’s t-test. **(C)** H&E-stained tissue cross-sections of organotypic cultures of NHEKs treated for 48 h with DMSO or 1 μM TG; inset shows separation between the basal and suprabasal layers in TG-treated cultures; scale bar = 100 μm. **(D)** Quantification of retained nuclei in cornified layers; data are shown as a box plot of the 25^th^-75^th^ percentile with a line at the median from N≥45 non-overlapping hpf from 3 biological replicates; P-value is from a two-tailed unpaired Student’s t test; (right) H&E-stained tissue cross-section from a TG-treated culture shows retention of nuclei in the cornified layers (magnified in inset); scale bar = 100 μm. **(E)** Immunostaining KRT10 and pERK in tissue cross-sections from organotypic cultures of NHEKs after 48 h of DMSO or TG treatment; scale bar = 50 µm; dashed line marks bottom of epidermis. (F) Quantification of tissue immunostaining of KRT10 (relative to Hoechst) and pERK (relative to total ERK) in organotypic cultures of NHEKs treated with DMSO or TG; data are shown as a box plot of the 25^th^-75^th^ percentile with a line at the median from N≥98 non-overlapping hpf per condition from 4 biological replicates; control mean normalized to 1; P-values are from two-tailed unpaired Student’s t-test.

Supporting our data from both RNA sequencing and proteomic analysis, TG-treated organotypic epidermis exhibited a reduction in KRT10 with immunostaining of tissue cross-sections demonstrating patchy loss of this cytoskeletal element in the suprabasal layers (Fig. 4E-F, upper panels). Disruption of KRT10 function can compromise epidermal integrity and cause abnormal cornification as in patients with *KRT10* mutations in epidermolytic ichthyosis (48, 58, 59). Like in SERCA2-deficient keratinocytes, ERK hyper-activation was also seen upon TG treatment of NHEK epidermal cultures; pERK was highest in the lower keratinocyte layers, where intercellular adhesion was most compromised (Fig. 4E-F, lower panels). These findings supported our hypothesis that loss of SERCA2 function disrupted tissue integrity and epidermal differentiation though over-activation of the MAPK pathway via ERK, resulting in depletion of KRT10.

### Biopsies of Darier disease exhibit mis-localization of cadherins, loss of keratin 10, and elevated ERK activation

To directly test if our *in vitro* data from SERCA2-depleted or –inhibited epidermal cultures were reflective of DD pathogenesis, we examined de-identified skin biopsies from patients with DD. Similar to the pathologic features we found in organotypic cultures having SERCA2 genetically ablated or chemically inhibited, histologic examination of DD biopsies demonstrated loss of keratinocyte cohesion and defective differentiation with retention of nuclei in the cornified layers (Fig. 5A, inset). Immunostaining biopsies from five DD patients compared to normal control skin demonstrated a significant reduction in KRT10 and a concomitant increase in ERK activation within lesional skin (Fig. 5B), consistent with the findings from our *in vitro* human tissue model. Like in organotypic epidermis, analysis of immunostained DD biopsy cross-sections revealed a significant reduction in overall KRT10 intensity with patchy loss of the cytoskeletal element within DD lesions (Fig. 5C).

**Figure 5.**
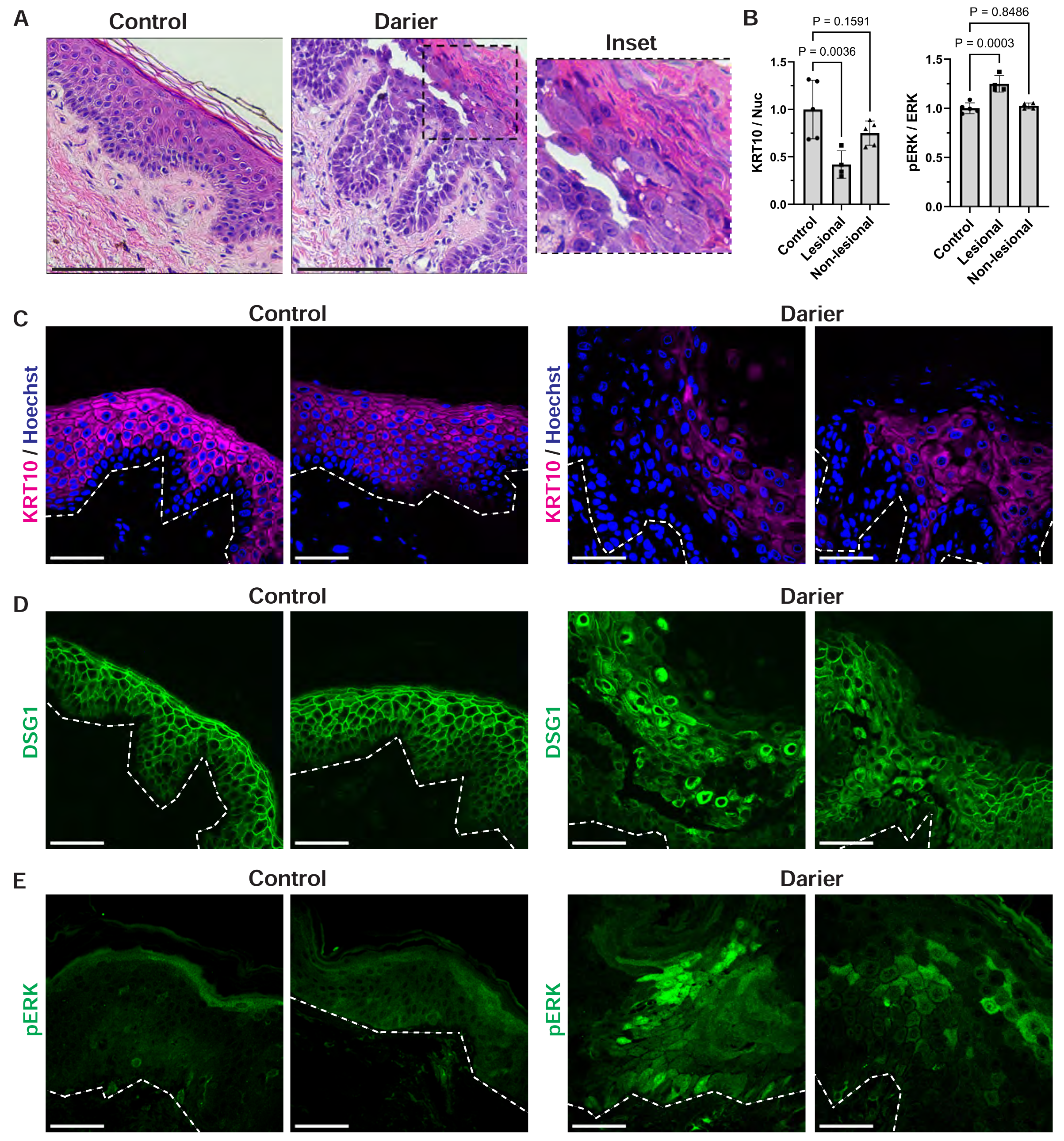
Biopsies of Darier disease exhibit mis-localization of cadherins, loss of keratin 10, and elevated ERK activation. **(A)** H&E-stained cross-sections of punch biopsies from the skin of control donors versus patients with Darier disease, which demonstrate loss of keratinocyte cohesion (acantholysis) and aberrant cornification (dyskeratosis) with retention of nuclei in the cornified layers (magnified in inset); scale bar = 50 µm. **(B)** Quantification of immunostaining of KRT10 (relative to Hoechst) and pERK (relative to total ERK) in cross-sections of control skin versus in lesional or non-lesional portions of Darier disease biopsies; bar graph depicts the mean ± SD from N≥20 non-overlapping hpf per group from 5 control and 5 Darier disease patients; control mean normalized to 1; P-values are from one-way ANOVA with Dunnett adjustment for multiple comparisons to control skin. **(C)** Immunostaining of KRT10, (**D**) DSG1, and (**E**) pERK in tissue cross-sections from patient biopsies; images shown are from 2 control and 2 Darier disease patients and are representative of N=5 patients in each group; scale bar = 50 µm; dashed line marks bottom of epidermis.

In addition, we found that protein levels of DSG1, the primary cadherin mediator of desmosomal adhesion in the upper epidermal layers (60, 61), were modestly reduced (data not shown). However, the localization of this cadherin was severely disrupted in DD lesions (Fig. 5D), collapsing around the nucleus instead of being concentrated at intercellular junctions. These findings agree with prior *in vitro* studies showing that keratinocytes having SERCA2 depleted by RNA interference retained normal levels of desmosomal components, but exhibited major defects in trafficking of these adhesive building blocks to the plasma membrane, thus impairing the cells’ ability to assemble strong intercellular junctions and compromising monolayer integrity (8, 26, 62, 63).

ERK activation assessed by immunostaining pERK, on the other hand, was significantly increased in biopsies from patients with DD compared to biopsies of normal skin (Fig. 5E). These data from DD tissue substantiate our findings from SERCA2-depleted or –inhibited human epidermal cultures and bolster our model in which hyper-activation of the MAPK pathway via ERK is a pathogenic driver in DD and represents a potential therapeutic target.

### Keratin expression and adhesive integrity in SERCA2-deficient keratinocytes are rescued by MEK inhibitors

We next assessed the effect of multiple selective MEK inhibitors on SERCA2-depleted keratinocytes to determine if elevated MEK activity directly contributes to the phenotype of SERCA2 deficiency. Quantification of keratinocyte mRNA levels by RT-PCR again confirmed that HET cells exhibit significantly reduced expression of the suprabasal keratins, KRT1 and KRT10, and DSG1 while the mRNA encoding KRT14 was not significantly different from controls (Fig. 6A). Importantly, treatment of HET cells with any of three MEK inhibitors (trametinib, U0126, PD184352) was sufficient to augment expression of KRT1, KRT10, and DSG1, while they did not significantly alter KRT14 mRNA levels.

**Figure 6.**
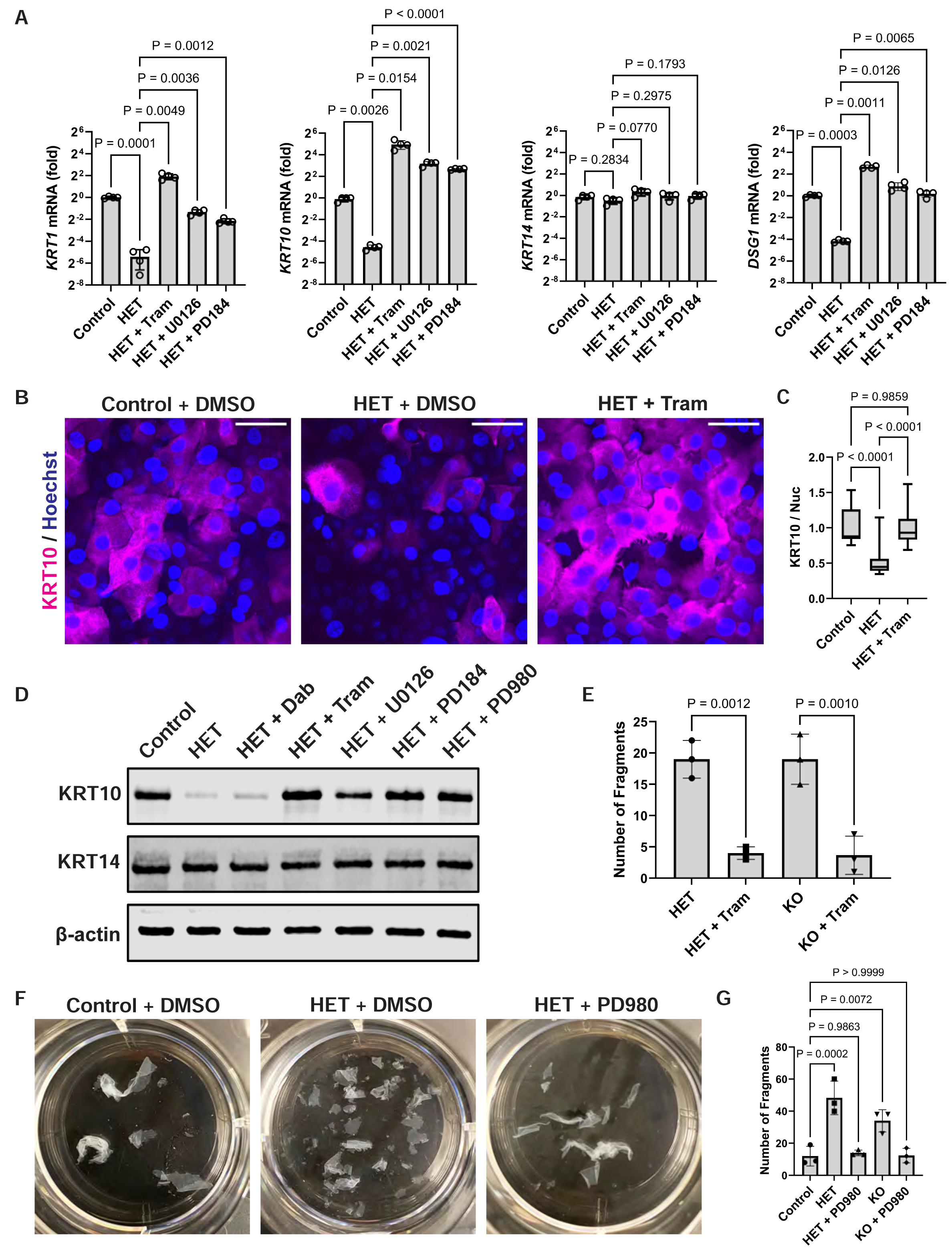
Keratin expression and adhesive integrity in SERCA2-deficient keratinocytes are rescued by MEK inhibitors. **(A)** Control or HET cells were differentiated in E-medium containing DMSO or the indicated MEK inhibitors for 24 h, then levels of the indicated transcripts were determined by RT-qPCR; data are graphed as the mean ± SD from N=4 biological replicates; P-values are from ANOVA (Brown-Forsythe) with Dunnett’s T3 method to adjust for multiple pair-wise comparisons. **(B)** Immunostaining of control or HET cells treated with DMSO or trametinib (Tram) at 1 μM for 48 h; images are representative of 2 independent experiments using 2 control and 2 SERCA2-deficient cell lines; scale bar = 50 µm**. (C)** Quantification of KRT10 immunostaining in control versus SERCA2-deficient cells treated with DMSO or Tram at 1 μM for 24 h; data are shown as a box plot of the 25^th^-75^th^ percentile with a line at the median from N≥21 non-overlapping hpf per group; control mean normalized to 1; P-values are from one-way ANOVA with Tukey adjustment for multiple pair-wise comparisons. **(D)** Immunoblot of KRT10 and KRT14 (β-actin is a loading control) in lysates from control versus HET cells treated with DMSO, dabrafenib (Dab) at 1 μM or MEK inhibitors (Tram at 1 μM; U0126 at 10 μM; PD184 at 10 μM; PD980 at 20 μM); data are representative of 2 independent experiments. **(E)** Quantification of epithelial fragments after mechanical stress was applied to HET and KO monolayers treated with DMSO or trametinib for 48 h; bar graphs display the mean ± SD with individual data points plotted for N=3 biological replicates; P-values are from one-way ANOVA with Tukey adjustment for multiple pair-wise comparisons. **(F)** Representative images of fragmented monolayers transferred into 6-well cell culture plates are shown for control and HET cells treated with DMSO or 20 μM PD980 for 24 h. (**G**) Quantification of epithelial fragments after mechanical stress was applied to control, HET, or KO monolayers treated with DMSO or PD980 at 20 μM for 24 h; bar graphs display the mean ± SD with individual data points plotted for N=3 biological replicates; P-values are from one-way ANOVA with Dunnett adjustment for multiple comparisons to control cells.

These differences in mRNA translated to changes in protein levels with HET cells exhibiting significantly reduced KRT10 as shown by immunostaining differentiated keratinocyte cultures (Fig. 6B). Consistent with its effect on *Krt10* mRNA, MEK inhibition increased KRT10 protein, making its levels comparable to that of control cells (Fig. 6C). Assessing keratinocyte lysates by immunoblot similarly demonstrated a reduction in KRT10, which multiple individual MEK inhibitors restored to near the baseline level of controls (Fig. 6D). In contrast, KRT14 levels were comparable in control and HET cells and were not appreciably altered by MEK inhibitors. Interestingly, targeting the MAPK pathway upstream of MEK using dabrafenib to inhibit RAF failed to rescue KRT10 expression. This result is consistent with prior data indicating that chronic suppression of RAF leads to paradoxical downstream *elevation* in MEK activity (64). This unintended result of RAF blockade changed the standard-of-care for patients with RAF-driven cancers to include dual RAF and MEK inhibitors to mitigate adverse effects of RAF inhibitors (54, 65). In fact, MEK hyper-activation in patients treated with RAF inhibitor monotherapy can cause a DD-like skin eruption with biopsies showing impaired keratinocyte adhesion and defective epidermal cornification (66–69).

Since trametinib robustly rescued the expression of adhesive proteins in HET cells and is an FDA-approved drug for the treatment of RAF-mutated cancers (70, 71), we investigated whether this compound could augment intercellular adhesion in SERCA2-deficient keratinocytes. Indeed, treatment with trametinib enhanced the resistance of HET or KO keratinocyte sheets to disruption by mechanical stress (Fig. 6E). An additional MEK inhibitor PD98059 similarly rescued cohesion of SERCA2-depleted cultures (Fig. 6F-G), further supporting the potential clinical utility of inhibiting this kinase to treat DD.

### MEK inhibitors promote keratinocyte cohesion and mitigate epidermal tissue disruption from SERCA2 inhibition

To confirm that ERK hyper-activation drives the pathogenic effects of reduced SERCA2 function, we also tested whether MEK inhibitors could reverse TG-induced loss of keratinocyte adhesion and impaired epidermal differentiation. Treatment of NHEKs with TG suppressed their ability to up-regulate KRT10 as indicated by immunostaining differentiated keratinocytes (Fig. 7A-B) and this effect of TG was mitigated by concomitant treatment with trametinib to inhibit MEK.

**Figure 7.**
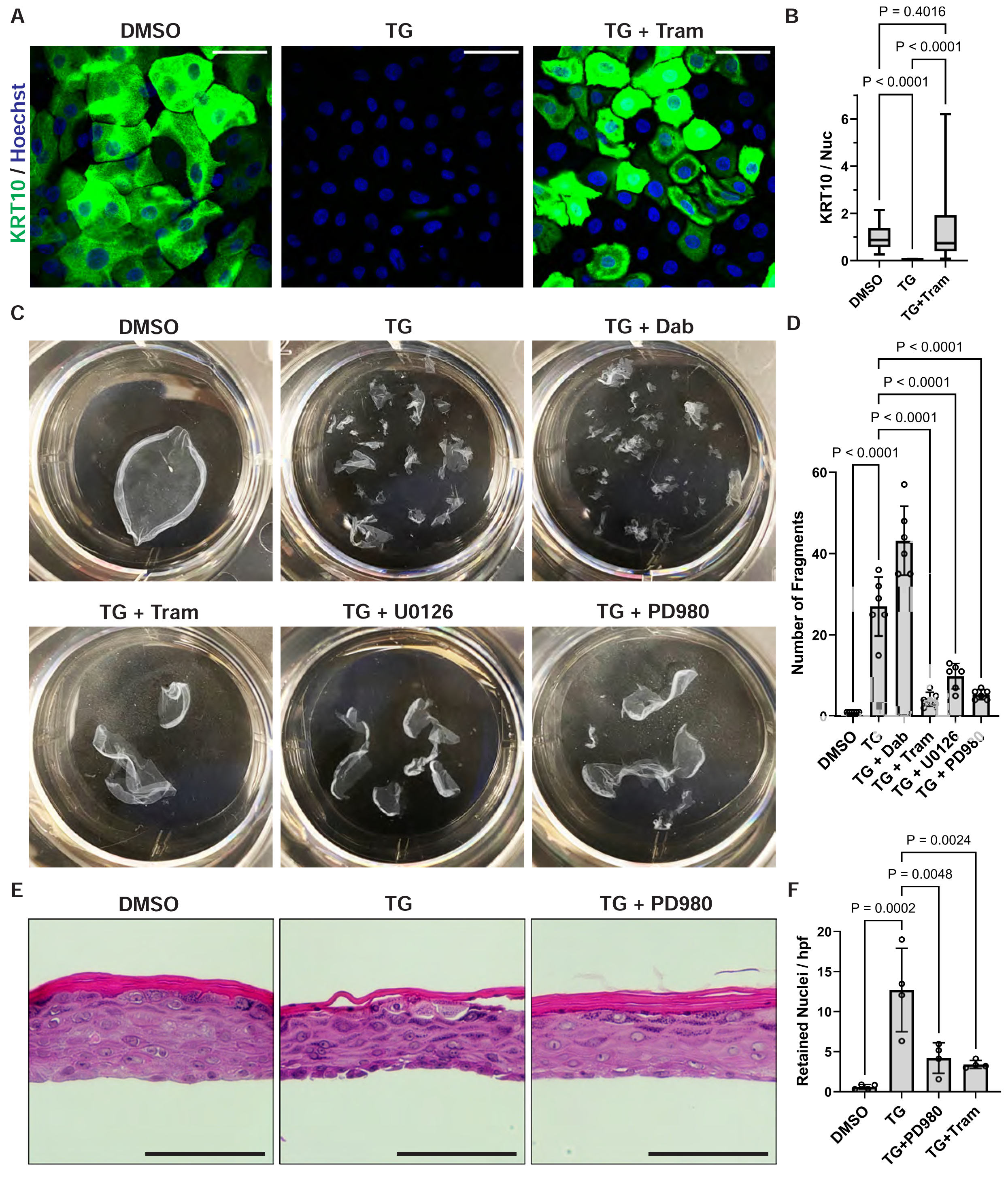
MEK inhibitors promote keratinocyte cohesion and mitigate epidermal tissue disruption from SERCA2 inhibition. **(A)** Immunostaining of KRT10 (and Hoechst nuclear stain) in THEKs treated with DMSO, 1 μM TG, or 1 μM TG plus 1 μM Tram (TG + Tram) for 48 h; scale bar = 50 µm. **(B)** Quantification of KRT10 immunostaining in THEKs treated with DMSO, TG, or TG plus Tram; data are shown as a box plot of the 25^th^-75^th^ percentile with a line at the median from N≥31 non-overlapping hpf from 2 independent THEK lines for both control and SERCA2-deficient cells; control mean normalized to 1; P-values are from one-way ANOVA with Tukey adjustment for multiple pair-wise comparisons. **(C)** Representative images of fragmented monolayers transferred into 6-well cell culture plates are shown for NHEKs treated with DMSO versus 1μM TG alone or with 1 μM dabrafenib versus a MEK inhibitor (1 μM Tram, 10 μM U0126, or 20 μM PD980) for 24h. (**D**) Quantification of epithelial fragments of NHEK monolayers; bar graphs display the mean ± SD with individual data points plotted for N=3 biological replicates; P-values are from one-way ANOVA with Tukey adjustment for multiple pair-wise comparisons. **(E)** H&E-stained tissue cross-sections of organotypic cultures of NHEKs treated with DMSO, 1 μM TG, or 1 μM TG plus 20 μM PD980, the latter displaying improved keratinocyte cohesion and normalization of cornification; scale bar = 100 µm. **(F)** Quantification of retained nuclei in cornified layers of organotypic cultures treated with the indicated inhibitors for 48 h; bar graph displays the mean ± SD with plotted individual values averaged from ≥49 non-overlapping hpf per condition from N=4 biological replicates; P-values are from one-way ANOVA with Tukey adjustment for multiple pair-wise comparisons.

This effect of MEK suppression on KRT10 in SERCA2-inhibited keratinocytes translated into a restoration of intercellular adhesive strength. While TG-treated NHEK monolayers readily fragmented upon mechanical stress, the integrity of epithelial sheets was restored by MEK inhibition with trametinib, U0126, or PD98059 despite SERCA2 inhibition (Fig. 7C-D). In contrast, upstream inhibition of RAF with dabrafenib did not promote intercellular adhesion, but rather exacerbated the effect of TG, further reducing the integrity of NHEK monolayers. These results are consistent with the reported ability of dabrafenib monotherapy to cause paradoxical hyper-activation of MEK (64), which we propose drives the pathogenic effects of SERCA2 deficiency and would explain the similar findings in biopsies of RAF inhibitor-induced Grover disease and DD.

We next treated mature organotypic epidermis with TG either alone or with PD98059 or trametinib to determine if MEK inhibition could prevent DD-like pathologic changes in organotypic epidermis. While TG-treated cultures exhibited loss of keratinocyte cohesion within the epidermal tissue, this effect was mitigated by MEK inhibitors (Fig. 7E). Moreover, treatment with PD98059 or trametinib reduced TG-induced retention of nuclei in the cornified layers (Fig. 7F), indicating that MEK inhibition could normalize epidermal differentiation in DD. Together, these data support our model in which the pathogenic effects of SERCA2 loss-of-function in epidermis are driven by ERK hyper-activation, which can be mitigated by MEK inhibition (see Graphical Abstract).

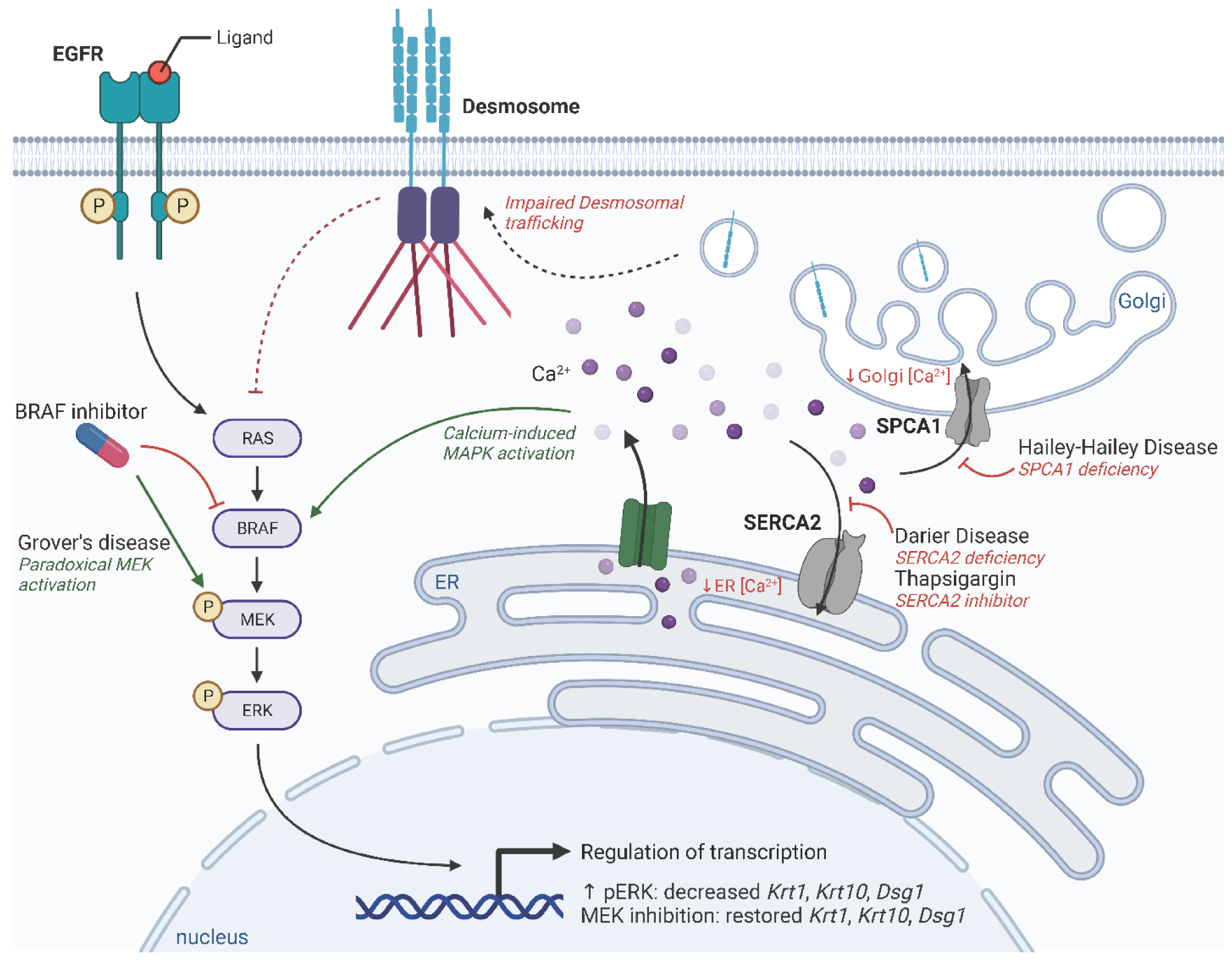
GRAPHICAL ABSTRACT.

## DISCUSSION

The advent of rational molecular therapeutics has revolutionized the treatment of many dermatologic conditions (4, 5); however, most advances have been made for common inflammatory diseases, like psoriasis and atopic dermatitis, driven by soluble cytokines amenable to targeting by monoclonal antibodies (72, 73). Unfortunately, skin blistering diseases and disorders of cornification are rare and most lack any FDA-approved treatments, though recent work suggests cytokine targeting may improve certain subtypes of ichthyosis (74–76). While we did not identify an inflammatory signature in our keratinocyte cultures, SERCA2-deficient cells had reduced IFI27 and other interferon-response genes, which might explain the susceptibility of DD lesions to viral infection (13). For DD, the knowledge of its genetic etiology has not significantly advanced the treatment of this chronic, incurable disorder. Though retinoids, which modulate keratinocyte differentiation, may be helpful in DD (14, 15), they are potent teratogens and can cause multi-system toxicity with long-term use (17, 18). Some evidence suggests reducing ER stress may be a viable alternative approach for DD treatment (9), though this strategy has not yet been tested clinically.

Using mice as a pre-clinical model has enabled important discoveries about disease pathogenesis that translated into new therapies for patients, but animal models do not always phenocopy human pathology, as with disrupting *Atp2a2* in mice (20, 21). The development of organoid models to replicate human tissues *in vitro* has the potential to revolutionize the way we investigate disease pathogenesis, identify lead compounds, and screen for therapeutic efficacy and toxicity (77, 78). Here, we leveraged gene editing in human keratinocytes and an organotypic epidermal model to elucidate the pathogenic mechanisms underlying a rare genetic disorder, which allowed us to identify much-needed potential therapeutic avenues. Though testing if our findings translate to an effective treatment approach for patients remains to be done, our work nevertheless underscores the power of organoids to generate large-scale transcriptomic and proteomic datasets that can be mined for rational drug targets to validate both *in vitro* and in patient-derived tissues (77, 79–82).

While the primary DD pathology derives from impaired the ER calcium homeostasis (26, 57), direct targeting of SERCA2 is likely to be therapeutically challenging. Recent intravital imaging of murine epidermis revealed that calcium flux undergoes dramatic shifts during epidermal proliferation (83) and in the late stages of cornification (30). Thus, sustained drug-induced activation of SERCA2 could compromise the dynamic nature of calcium-mediated signals critical for epidermal functions (27, 28). Since calcium is a major second messenger regulating diverse biological functions including skeletal muscle contraction, cardiac pacing, neurotransmitter release, and apoptosis in many cell types (84, 85), compounds affecting intracellular calcium may have a narrow therapeutic window. In fact, coincidence of neuropsychiatric and cardiac disease with DD (6, 86) implies a role beyond the skin for SERCA2 (87), including in higher-order cognition and behavior (88–90).

The study of rare diseases can yield unexpected findings that improve our understanding of fundamental biology or the pathology of more common disorders (91). While our study focused on DD, we found that MEK inhibition robustly increased expression of KRT1 and KRT10. Given that multiple MEK inhibitors are approved for clinical use (56, 70, 92), these could be evaluated for treating skin fragility disorders due to defective keratins like epidermolytic ichthyosis and epidermolysis bullosa simplex, linked to *KRT1/KRT10* and *KRT5/KRT14* mutation*, respectively* (59, 93, 94). Since MEK inhibitors augment expression of wild-type KRT1 and KRT10 protein, these compounds might rescue keratinocyte integrity in these orphan diseases. Finally, our results demonstrating ERK hyper-activation in DD bring to mind a non-inherited dermatologic disorder called Grover disease, which can be induced by RAF inhibition (66, 67, 69). Given that biopsy findings in Grover disease can be indistinguishable from DD (55), it seems likely that they share a common pathogenic mechanism. The fact that RAF inhibitor-induced Grover disease is suppressed by adding a MEK inhibitor (54, 65) further supports our hypothesis that MEK inhibitors would be effective for DD.

In sum, our findings indicate that excess ERK signaling disrupts the differentiation and cohesion of epidermal keratinocytes (37, 39, 40, 51) to drive the pathogenesis of DD, which can be reversed by inhibiting MAPK signaling through MEK. Whether this pathway could be similarly targeted for therapeutic benefit in other diseases compromising desmosomal adhesion remains to be tested, but extensive data previously implicated a pathogenic role for the MAPK pathway in pemphigus via p38 (95–97), which unfortunately did not progress to clinical approval (98). However, given that multiple MEK inhibitors are already approved for clinical use with long-term data supporting their safety (56, 65, 70, 92), we propose that MEK inhibition could represent a viable therapeutic strategy for both inherited and acquired blistering diseases, skin fragility syndromes, and disorders of cornification, all of which are in great need of novel treatments.

## METHODS

### Reagents

The SERCA2 inhibitor thapsigargin (Cat. #12758), MEK inhibitors trametinib (Cat. #62206), U0126 (Cat. #9903), PD98059 (Cat. #9900), and PD184352 (Cat. #12147), and B-Raf inhibitor dabrafenib (Cat. #91942) are from Cell Signaling Technology. Rabbit antibodies against SERCA2 (D51B11; Cat. #9580), phospho-ERK1/2 (D13.14.4E; Cat. #4370) and mouse antibodies again ERK1/2 (L34F12; Cat. #4696) and β-Actin (8H10D10; Cat. #3700) are from Cell Signaling Technology. Rabbit anti-Cytokeratin 10 (Cat. #ab76318), mouse anti-Cytokeratin 14 (Cat. #ab7800), mouse anti-Desmoglein 1 (Cat. #ab12077), and mouse anti-Involucrin (Cat. #ab68) are from Abcam. For fluorescent immunoblotting, IRDye 800CW goat anti-rabbit IgG (Cat. #926-32211) and IRDye 680RD goat anti-mouse IgG (Cat. #926-68070) are from LI-COR Biosciences. For fluorescent immunostaining, secondary antibodies are from Thermo-Fisher: Goat anti-mouse IgG with Alexa Fluor 405 (Cat. #A31553), Alexa Fluor 488 (Cat. #A11001), Alexa Fluor 594 (Cat. #A11005), or Alexa Fluor 633 (Cat. #A21050); goat anti-rabbit IgG with Alexa Fluor 405 (Cat. #A31556), Alexa Fluor 488 (Cat. #A11008), Alexa Fluor 594 (Cat. #A11012), or Alexa Fluor 633 (Cat. #A21070). Hoechst 33342 is from Thermo-Fisher (Cat. #H1399).

### Cell culture

hTERT-immortalized human epidermal keratinocytes (THEKs) from original line N/TERT-2G (31) were grown in keratinocyte serum-free medium (KSFM) ordered as a kit (Thermo-Fisher Cat. # 37010022) with supplements of 0.2 ng/mL human epidermal growth factor and 30 µg/mL bovine pituitary extract. Other additives included 0.31 mM CaCl_2_, 100 U/mL penicillin, and 100 μg/mL streptomycin.

Normal human epidermal keratinocytes (NHEKs) were grown in Medium 154 with 0.07 mM CaCl_2_ (Thermo-Fisher Cat. #M154CF500) and 1x human keratinocyte growth supplement (Thermo-Fisher Cat. #S0015), and 1x gentamicin/amphotericin (Thermo-Fisher Cat. #R01510).

J2-3T3 immortalized murine fibroblasts were grown in complete Dulbecco’s Modified Eagle Medium (DMEM) (Thermo-Fisher Cat. #11965092) supplemented with 10% FBS (Hyclone, Fisher Scientific Cat. #SH3039603), 2 mM GlutaMAX (Thermo-Fisher Cat. #35050061), 100 U/mL penicillin, and 100 μg/mL streptomycin.

All cell lines were maintained at 37°C in 5% CO_2_ in an air-jacketed, humidified incubator. Cells were grown on sterile cell culture dishes and passaged at sub-confluency using 0.25% Trypsin-EDTA (Thermo-Fisher Cat. #15400054).

### Organotypic epidermal culture

Human organotypic epidermal “raft cultures” were generated as described (22, 99). Cultures were differentiated using E-medium, a 3:1 mixture of DMEM:Ham’s F12 (Thermo-Fisher Cat. #11765054) with 10% FBS, 180 µM adenine (Sigma Cat. #A2786), 0.4 µg/mL hydrocortisone (Sigma, Cat. #H0888), 5 µg/mL human insulin (Sigma Cat. #91077C), 0.1 nM cholera toxin (Sigma, Cat. #C8052), 5 µg/mL apo-transferrin (Sigma Cat. #T1147), and 1.36 ng/mL 3,3′,5-tri-iodo-L-thyronine (Sigma Cat. #T6397).

J2-3T3 fibroblasts were seeded into collagen matrix rafts within transwells (Corning Cat. #353091). For each raft, 1 x 10^6^ fibroblasts were resuspended in 1/10 the final volume of sterile filtered reconstitution buffer (1.1 g of NaHCO_3_ plus 2.39 g of HEPES in 50 mL 0.05 N NaOH), then 1/10 the final volume of 10X DMEM (Sigma-Aldrich Cat. #D2429) was added. The cells were mixed thoroughly by pipetting, then high-concentration rat tail collagen I (Corning Cat. #CB354249) was added (4 mg/mL final concentration), along with sterile diH_2_O to bring the solution up to the final volume. If necessary, 0.05 N NaOH was added dropwise to adjust the pH to ∼7 based on the phenol red indicator. The collagen-fibroblast slurry was mixed via inversion, then pipetted into a transwell insert placed within a deep 6-well cell culture plates (Corning Cat. #08-774-183). The rafts were polymerized at 37°C for one hour, after which they were submerged in complete DMEM and incubated at 37°C overnight.

Next, confluent keratinocyte cultures were trypsinized and resuspended in E-medium with EGF (5 ng/ml) to final concentration of 0.5 x 10^6^ cells/mL (2 mL per organotypic culture). The DMEM was aspirated from the upper and lower transwell chambers, then 2 mL (1 x 10^6^ cells) of the keratinocyte suspension were pipetted on top each raft. E-medium with EGF was added to the bottom transwell chamber to submerge the raft, and the cultures were incubated at 37°C. After 24 hours, E-medium was aspirated from the top and bottom chambers. An air-liquid interface was established to induce stratification by adding E-medium (without EGF) only to the bottom chamber to reach the bottom of the raft. Organotypic cultures were grown at 37°C for up to 12 days. E-medium in the bottom chamber was replaced every other day. For drug treatments, inhibitors or vehicle control (DMSO) were diluted in E-medium in the bottom chamber. Inhibitors were used at the following concentrations: Thapsigargin (1 µM), trametinib (1 µM), and PD98059 (20 µM). For histologic examination, the transwell was moved to a standard 6-well cell culture plate and submerged in 10% neutral-buffered formalin (Fisher Scientific Cat. #22-026-435) for at least 24h. Organotypic cultures were processed for histologic examination by Core A of the Penn SBDRC or the Experimental Histopathology Core of the Fred Hutchinson Cancer Center.

### CRISPR/Cas9 gene editing

CRISPR knockout (KO) keratinocytes were generated as described (24). Single-guide RNAs (sgRNAs) were designed to target the *ATP2A2* gene (sgRNA2: GTTTTGGCTTGGTTTGAAGA) or the *TUBAP* pseudogene (sgRNA1: GTATTCCGTGGGTGAACGGG) to generate a control KO line using a web tool for CRISPR design (https://portals.broadinstitute.org/gpp/public/analysis-tools/sgrna-design). Synthetic sgRNA target sequences were inserted into a cloning backbone, pSpCas9 (BB)-2A-GFP (PX458) (Addgene, Cat. #48138), and then cloned into competent *E. coli* (Thermo-Fisher, Cat. #C737303). Proper insertion was validated by Sanger sequencing. The final plasmid was transfected into an hTERT-immortalized human epidermal keratinocyte line (N/TERT-2G (31)) using the TransfeX transfection kit (ATCC Cat. #ACS4005) in the presence or absence of JAK1/JAK2 inhibitor, baricitinib (10 µg/mL). GFP-positive single cells were plated and expanded. Cells were genotyped and analyzed by Sanger sequencing to confirm the presence of heterozygous or homozygous mutations in the target gene.

### RNA sequencing (RNAseq)

RNAseq libraries were prepared for transcriptomics analysis as described (100). THEKs were grown to confluency in 10 cm cell culture dishes in KSFM, then differentiated in E-medium for 72 h. RNA was isolated from THEKs using the RNeasy kit (Qiagen) according to the manufacturer’s instructions. mRNAs were isolated using the NEBNext Poly(A) mRNA magnetic isolation module (New England Biolabs). RNAseq libraries were prepared for Illumina using the NEBNext Ultra-Directional RNA library preparation kit (New England Biolabs). Library quality was confirmed using an Agilent BioAnalyzer 2100 and quantified using the NEBNext Library Quant Kit for Illumina (New England Biolabs). Sequencing was performed using the Illumina NextSeq500 platform employing a single-end, 75-base pair sequencing strategy. All RNA-seq reads were then aligned to the *Homo sapiens* reference genome (UCSC hg19, RefSeq and Gencode gene annotations) using RNA STAR under default settings. Fragments per kilobase per million mapped fragments (FKPM) generation and differential expression analysis were done using the DESeq2 package. Statistical significance was determined using an adjusted *P* value (*P*_adj_) of ≤ 0.05.

### Gene Ontology Analyses

Gene ontology (GO) analyses of RNAseq data were performed as described (100) using PANTHER at http://pantherdb.org/. Enrichment analysis under the category “biological process” was performed to identify statistically overrepresented GO terms. Statistical significance was determined using Fisher’s exact test. Normalized enrichment scores and false discovery rates were computed for each biological process. GO terms were plotted using Prism 9.

### Whole-cell proteomics

THEKs were grown to confluency in 10 cm cell culture dishes in complete KSFM, then differentiated in E-medium for 72 h. Cells were then washed with PBS and lysed in ice-cold lysis buffer [0.1 M Ammonium Bicarbonate, 8 M Urea, 0.1% (w/v) RapiGest SF (Waters)]. Lysates were homogenized using a microtip probe sonicator (Fisher Scientific) and clarified by centrifugation. Clarified supernatants were transferred to a new microcentrifuge tube and protein concentrations were determined by BCA assay (Pierce). After normalizing sample volumes, 200 µg of protein from each sample were reduced by the addition of TCEP to a final concentration of 5 mM for 1 h at room temperature. Protein samples were then alkylated for 30 minutes at room temperature by adding iodoacetamide to a final concentration of 10 mM. Unreacted iodoacetamide was quenched by adding TCEP to a final concentration of 10 mM for 30 minutes at room temperature. Following reduction and alkylation, samples were diluted with 0.1 M ammonium bicarbonate to decrease the urea concentration to 1.5 M. Protein samples were then digested overnight at 37°C with Trypsin Gold (Promega) according to the manufacturer’s instructions. On the following day, each sample was adjusted to pH 2 with 2 M HCl to hydrolyze the RapiGest surfactant. The precipitated RapiGest was then removed via centrifugation at 10,000 x g for 5 minutes. Acetonitrile and trifluoroacetic acid were added to the clarified supernatants to final concentrations of 5% (v/v) and 0.1% (v/v), respectively. Trypsin-digested peptides were then desalted using MacroSpin C18 columns (Pierce), according to the manufacturer’s instructions. Following elution, peptides were completely dried under vacuum and resuspended in 0.1% (v/v) formic acid to a final concentration of ∼0.5 µg/mL. Samples were transferred into autosampler vials and subjected to mass spectrometry analysis using an Orbitrap Eclipse mass spectrometer (Thermo Scientific) as described (101). Raw spectral data were processed in MaxQuant using the *Homo sapiens* reference protein library (UniProt) and relative protein abundances were determined using the label-free quantification (LFQ) intensity method as described (101).

### RNA isolation and real-time quantitative PCR

THEKs were seeded at a density of 1 x 10^6^ cells per well of 6-well cell culture dishes in complete KSFM. Upon reaching confluence, KSFM was replaced with E-medium containing vehicle control (DMSO) or inhibitors at the following concentrations: Trametinib (1 µM), U0126 (10 µM), or PD184352 (10 µM). At the indicated time points, cell culture medium was removed and cells were washed once with PBS. RNA was extracted using the NucleoSpin RNA Plus Kit (Machery-Nagel) followed by reverse-transcription with the iScript cDNA synthesis kit (Bio-Rad) according to the manufacturer’s instructions. cDNAs were diluted 5-fold and 2 µL of the resulting cDNAs were used for each 10 µL qPCR assay. Real-time qPCR assays were carried out using the TaqMan Gene expression master mix (Applied Biosystems) according to the manufacturer’s protocol. Pre-designed TaqMan probes from Thermo-Fisher were used for quantification of human *ACTB* (Hs01060665_g1), *KRT1* (Hs00196158_m1), *KRT10* (Hs00166289_m1), *KRT14* (Hs00265033_m1), *DSG1* (Hs00355084_m1), *ALOX12B* (Hs00153961_m1), *ALOX15B* (Hs00153988_m1), *IFI27* (Hs01086373_g1), *TRPV3* (Hs00376854_m1), *TRPV4* (Hs01099348_m1), or *NOTCH1* (Hs01062014_m1). The housekeeping gene *ACTB* was used as an endogenous control and mRNA fold changes were determined using the 2^(-ΔΔCt)^ method.

### Immunoblotting

THEKs were seeded at a density of 1 x 10^6^ cells per well of 6-well cell culture dishes in KSFM. Upon reaching confluence, KSFM was replaced with E-medium containing vehicle control (DMSO) or inhibitors at the following concentrations: Dabrafenib (1 µM), trametinib (1 µM), U0126 (10 µM), PD184352 (10 µM), or PD98059 (20 µM). After 72 h, whole-cell lysates were generated by washing cells once in PBS followed by lysis in urea sample buffer [8 M Urea, 60 mM Tris, 1% SDS, 10% glycerol, 5% β-mercaptoethanol, 0.0005% pyronin-Y, pH 6.8] for 10 min. Lysates were homogenized using a microtip probe sonicator (Fisher Scientific).

Whole-cell lysates were loaded onto Any kD Mini-PROTEAN TGX Precast Protein Gels (Bio-Rad) and separated by electrophoresis. Proteins were transferred onto nitrocellulose membranes (Bio-Rad) in ice-cold Towbin transfer buffer (25 mM Tris, 192 mM glycine, 20% (v/v) methanol) at 100 V for 90 minutes at 4°C. Membranes were blocked in Intercept tris-buffered saline (TBS) blocking buffer (LI-COR) for 45 min at room temperature. Membranes were probed overnight at 4°C with the primary antibodies diluted at 1:1000 in Intercept TBS blocking buffer (LI-COR). Blots were washed at least three times in 1x TBS containing 0.1% (v/v) Tween-20 (TBS-T), then incubated for 1 h at room temperature with IRDye 800CW goat anti-rabbit IgG and/or IRDye 680RD goat anti-mouse IgG (LI-COR) diluted 1:10,000 in Intercept TBS blocking buffer. Blots were washed at least three times in TBS-T and proteins were visualized using an Odyssey Fc Imaging System (LI-COR).

### Fluorescent immunocytochemistry

Keratinocytes were grown to confluency in 35 mm glass-bottom cell culture dishes (MatTek #P35G-1.5-20-C). For staining keratins and desmosomal proteins, cells were fixed in ice-cold 100% methanol at -20°C for 2 min, allowed to dry, then re-hydrated in PBS. For staining other proteins, cells were fixed in 4% paraformaldehyde for 10 min at 37°C. Fixed cells were then incubated with blocking buffer [0.5% (w/v) bovine serum albumin (BSA, Sigma), 10% (w/v) normal goat serum (NGS, Sigma) in PBS] for 30 min at 37°C. Cells were then washed with PBS. Primary antibodies were diluted in 0.5% (w/v) BSA in PBS and incubated on the cells overnight at 4°C. Primary antibody dilutions were as follows: rabbit anti-cytokeratin 10 (1:2000) and rabbit anti-SERCA2 (1:200). Cells were washed three times with PBS, then incubated with species-specific secondary antibodies diluted at 1:300 (with or without Hoechst at 1:500) in 0.5% (w/v) BSA in PBS for 30 min at 37°C. Cells were washed three times with PBS, then held in PBS for imaging using spinning-disk confocal microscopy, as detailed below.

### Histologic analysis and tissue procurement

Paraffin-embedded formalin-fixed tissue cross-sections of organotypic epidermis or skin biopsies were processed for histology and stained with hematoxylin and eosin (H&E) using standard methods. H&E-stained glass slides were imaged on an EVOS FL imaging system (Thermo-Fisher) using an EVOS 40X long working distance, achromatic, phase-contrast objective (Thermo-Fisher). Images were captured using the embedded high-sensitivity interline CCD color camera.

### Fluorescent immunohistochemistry

Paraffin-embedded formalin-fixed tissue cross-sections on glass slides were incubated at 65°C for 2 h. Sections were prepared for staining by immersion in 3 baths of xylenes (Fisher) for 5 min each, followed by 3 baths of 95% ethanol for 5 min each, then 70% ethanol for 5 minutes, and finally 3 baths of PBS for 5 min each. Slides were then submerged in antigen retrieval solution [0.1 M sodium citrate (pH 6.0) with 0.05% (v/v) Tween-20] and heated to 95°C for 15 min. Slides were allowed to cool to room temperature and then washed with PBS. Tissue sections were encircled with a hydrophobic barrier using a PAP pen. Tissue sections were incubated in blocking buffer [0.5% (w/v) BSA, 10% (v/v) NGS in PBS] for 30 min at 37°C in a humidified chamber. Slides were washed in 3 baths of PBS for 5 min each, then incubated with primary antibodies diluted in 0.5% (w/v) BSA in PBS overnight at 4°C in a humidified chamber. Primary antibody dilutions were as follows: rabbit anti-cytokeratin 10 (1:3000), mouse anti-desmoglein 1 (1:50), rabbit anti-phospho-ERK (1:400), mouse anti-ERK (1:400). Slides were then washed in 3 baths of PBS for 5 min each and incubated with secondary antibodies diluted at 1:300 (with or without Hoechst at 1:500) in 0.5% (w/v) BSA in PBS for 60 min at 37°C in a humidified chamber. Slides were washed in 3 baths of PBS for 5 min each and mounted in Prolong Gold (Invitrogen Cat. #P36934) with a glass coverslip applied over the tissue sections. Slides were allowed to dry overnight prior to imaging by spinning-disk confocal microscopy, as detailed below.

### Fluorescence Microscopy Imaging

Images were acquired on a Hamamatsu ORCA-FusionBT sCMOS camera using a Yokogawa W1 spinning-disk confocal (SDC) system on a Nikon Ti2 microscope. Samples were illuminated using 405, 488, 561, and 640 nm laser excitation lines and fluorescence was detected using a 60x 1.2 NA water objective (Nikon) with standard emission filters.

For live imaging of cytoplasmic calcium, THEKs were transduced with GCaMP6 (Addgene Cat. # 40753) sub-cloned into the pLZRS retroviral vector. HEK293 Phoenix cells were grown in complete DMEM, then transfected with 4 μg pLZRS-GCaMP6 DNA plus 12 μl FuGENE 6 (Promega Cat. #E2691) in 800 μl of Opti-MEM (Thermo-Fisher Cat. #31985070), which was added to the cells and left overnight. Retroviral supernatants were collected the next day and polybrene (Sigma Cat. #H9268) was added at a concentration of 4 μg/ml. KSFM was removed from THEKs and replaced with viral medium for 1 h at 37°C. Cells were then washed in PBS and placed back in their normal medium and expanded in culture.

GCaMP6-transduced cells were seeded into 35 mm glass-bottom dishes in low-calcium medium (0.31 mM) and grown to confluency, then were imaged by confocal microscopy at 1 frame every 5 s. Cells were exposed to high-calcium (1.3 mM) after 60 s, then were imaged for an additional 5 min at 1 frame every 5 s. The Fiji “Measure” function was used to calculate the mean intensity of GCaMP6 signal across the entire high-powered field (hpf) of the time series of images. The intensity at 0 s was subtracted from the intensity at each subsequent time point to calculate the change from baseline as a function of time across independent replicates for each genotype.

### Fluorescent Microscopy Quantification

Fluorescence microscopy images were analyzed using Fiji software. Quantification of fluorescence intensity was performed in a blinded manner using non-visibly identifiable microscopy images. The Fiji “Measure” function was used to calculate the mean intensity across the entire hpf of fixed cells or a region of interest (ROI) encompassing the entire epithelium circumscribed using the polygon selection tool. The mean intensity for each condition was averaged across multiple independent hpf for each biological replicate.

### Tissue Morphologic Quantification

Counts of retained nuclei in the cornified layers were performed by hand using non-visibly labeled H&E images to identify and count the numbers of retained nuclei per non-overlapping hpf for each condition, which were averaged across multiple independent experiments.

### Fluorescent Tissue Staining Quantification

Fluorescence images of immunostained tissue sections were captured by SDC microscopy as above and analyzed using Fiji. Quantification of fluorescence intensity was performed in using non-visibly labeled images from immunostained tissue cross-sections. The Fiji “Measure” function was used to calculate the mean intensity of the fluorescence signal and the mean background fluorescence intensity, which was subtracted from each value. The net fluorescence intensity was averaged across multiple non-overlapping hpf and the values from control cultures were normalized to an average value of 1.

### Monolayer Mechanical Dissociation Assay

Dispase-based mechanical dissociation assays were carried out as described (102). Keratinocytes were plated at a density of 1 x 10^6^ cells per well of 6-well cell culture dishes. Upon reaching confluence, the calcium concentration of the medium was adjusted to 1.3 mM. Vehicle control (DMSO) or chemical inhibitors were added at the following concentrations: Thapsigargin (1 µM), dabrafenib (1 µM), trametinib (1 µM), U0126 (10 µM), PD184352 (10 µM), or PD98059 (20 µM). After 24 to 72 h, monolayers were washed with PBS and then incubated with 500 μL dispase (5 U/ml) in Hank’s balanced salt solution (Stemcell Technologies, Cat. #07913) for 30 min at 37°C. Next, 4.5 ml PBS was added to the wells and all liquid plus released monolayers were transferred into 15 mL conical tubes, which were placed together in a rack and inverted 5-10 times to induce mechanical stress. Monolayer fragments were transferred back into 6-well cell culture plates and imaged with a 12-megapixel digital camera. Fragments were counted in Fiji.

### Statistics

Statistical analyses were performed using the open-source statistics package R or Prism version 9 (GraphPad), which was used to generate graphs. Statistical parameters including sample size, definition of center, dispersion measures, and statistical tests are included in each figure legend. Datasets were tested for normality using the D’Agostino-Pearson test. The means of two normally distributed groups were compared using a two-tailed unpaired Student’s t test. The means of more than two normally distributed groups were compared using a one-way ordinary ANOVA followed by P-value adjustment for multiple comparisons. P-values less than 0.05 were considered statistically significant. Exact P-values are included in each figure.

### Study Approval

Normal human epidermal keratinocytes (NHEKs) from de-identified neonatal foreskins were procured by the Penn Skin Biology and Diseases Resource-based Center (SBDRC) under a protocol (#808224) approved by the University of Pennsylvania Institutional Review Board (IRB). Tissue cross-sections of de-identified skin biopsies were obtained from a tissue bank by the Penn SBDRC under a protocol (#808225) approved by the University of Pennsylvania IRB. The use of de-identified tissues collected for clinical purposes that would otherwise be discarded was deemed exempt by the IRB for written informed consent.

## AUTHOR CONTRIBUTIONS

Conceptualization: S.A.Z. and C.L.S. Data Curation: S.A.Z., S.E., J.Z., A.T., B.C.C., and C.L.S. Formal analysis: S.A.Z., S.E., J.Z., A.T., B.C.C., C.L.S. Investigation: S.A.Z., M.K.S., S.E., J.Z., A.T., and C.L.S. Resources: B.C.C., J.E.G, and C.L.S. Supervision: C.L.S. Visualization: S.A.Z., S.E., and C.L.S. Validation: S.A.Z. and C.L.S. Writing (original draft): S.A.Z. and C.L.S. Writing (review and editing): S.A.Z., M.K.S., S.E., J.Z. A.T., B.C.C., J.E.G., and C.L.S.

## ACKNOWLEDGEMENTS

## Acknowledgements

The authors would like to thank members of the Simpson Lab for helpful discussions and critical reading of the manuscript. We thank the Penn Skin Biology and Diseases Resource-Based center (NIH P30-AR069589) for supporting this work with histology services and archived biopsy specimens (Stephen Prouty) and NHEK isolation (Christine Marshall). We thank the Fred Hutchinson Cancer Center Experimental Histopathology Core for histology services. We thank Dr. Andi Liu and Dr. Joseph Mougous (University of Washington) for assistance with whole-cell proteomics. We thank Dr. Joshua Woodward (University of Washington) for access to reagents and equipment. S.A.Z. was supported by the Cora May Poncin fellowship fund. M.K.S. and J.E.G. were supported by NIH P30-AR075043, R21-AR077741 and National Psoriasis Foundation Translational Research Grant 852098. S.E. was supported by NIH T32-AR007465 and F99-CA264315. B.C.C. was supported by NIH R01-AR077615. C.L.S. was supported by NIH K08-AR075846 as well as an Innovation Pilot Award and Genomics Core Pilot Award through the University of Washington Institute for Stem Cell and Regenerative Medicine. Graphical abstract was generated using BioRender.com.

## REFERENCES

1. Boyden LM, Choate KA. The Molecular Revolution in Cutaneous Biology: Identification of Skin Disease Genes. J Invest Dermatol. 2017; 137(5): p. e61–e65.

2. Sarig O, Sprecher E. The Molecular Revolution in Cutaneous Biology: Era of Next-Generation Sequencing. J Invest Dermatol. 2017; 137(5): p. e79–e82.

3. Salik D, et al. Clinical and molecular diagnosis of genodermatoses: Review and perspectives. J Eur Acad Dermatol Venereol. 2023; 37(3): p. 488–500.

4. Silverberg N. Emerging therapies in genodermatoses. Clin Dermatol. 2020; 38(4): p. 462–466.

5. Hill CR, Theos A. What’s New in Genetic Skin Diseases. Dermatol Clin. 2019; 37(2): p. 229–239.

6. Jacobsen NJ, et al. ATP2A2 mutations in Darier’s disease and their relationship to neuropsychiatric phenotypes. Hum Mol Genet. 1999; 8(9): p. 1631–6.

7. Sakuntabhai A, et al. Mutations in ATP2A2, encoding a Ca2+ pump, cause Darier disease. Nat Genet. 1999; 21(3): p. 271–7.

8. Hobbs RP, et al. The calcium ATPase SERCA2 regulates desmoplakin dynamics and intercellular adhesive strength through modulation of PKCα signaling. FASEB J. 2011; 25(3): p. 990–1001.

9. Savignac M, et al. SERCA2 dysfunction in Darier disease causes endoplasmic reticulum stress and impaired cell-to-cell adhesion strength: rescue by Miglustat. J Invest Dermatol. 2014; 134(7): p. 1961–1970.

10. Takagi A, et al. Darier disease. J Dermatol. 2016; 43(3): p. 275–9.

11. Nellen RG, et al. Mendelian Disorders of Cornification Caused by Defects in Intracellular Calcium Pumps: Mutation Update and Database for Variants in ATP2A2 and ATP2C1 Associated with Darier Disease and Hailey-Hailey Disease. Hum Mutat. 2017; 38(4): p. 343–356.

12. Dodiuk-Gad R, et al. Bacteriological aspects of Darier’s disease. J Eur Acad Dermatol Venereol. 2013; 27(11): p. 1405–9.

13. Vogt KA, et al. Kaposi varicelliform eruption in patients with Darier disease: a 20-year retrospective study. J Am Acad Dermatol. 2015; 72(3): p. 481–4.

14. Hanna N, et al. Therapeutic Options for the Treatment of Darier’s Disease: A Comprehensive Review of the Literature. J Cutan Med Surg. 2022; 26(3): p. 280–290.

15. Christophersen J, et al. A double-blind comparison of acitretin and etretinate in the treatment of Darier’s disease. Acta Derm Venereol. 1992; 72(2): p. 150–2.

16. Dicken CH, et al. Isotretinoin treatment of Darier’s disease. J Am Acad Dermatol. 1982; 6(4 Pt 2 Suppl): p. 721–6.

17. Zaenglein AL, et al. Consensus recommendations for the use of retinoids in ichthyosis and other disorders of cornification in children and adolescents. Pediatr Dermatol. 2021; 38(1): p. 164–180.

18. Doolan BJ, et al. Retinoid-induced skeletal hyperostosis in disorders of keratinization. Clin Exp Dermatol. 2022; 47(12): p. 2273–2276.

19. Ashok Kumar P, et al. Debilitating Darier’s Disease and Its Impact on the Quality of Life. Cureus. 2020; 12(5): p. e8133.

20. Prasad V, et al. Haploinsufficiency of Atp2a2, encoding the sarco(endo)plasmic reticulum Ca2+-ATPase isoform 2 Ca2+ pump, predisposes mice to squamous cell tumors via a novel mode of cancer susceptibility. Cancer Res. 2005; 65(19): p. 8655–61.

21. Toki H, et al. Novel allelic mutations in murine Serca2 induce differential development of squamous cell tumors. Biochem Biophys Res Commun. 2016; 476(4): p. 175–182.

22. Simpson CL, et al. RNA interference in keratinocytes and an organotypic model of human epidermis. Methods Mol Biol. 2010; 585: p. 127–46.

23. Guitart JR, Jr., et al. Research Techniques Made Simple: The Application of CRISPR-Cas9 and Genome Editing in Investigative Dermatology. J Invest Dermatol. 2016; 136(9): p. e87–e93.

24. Sarkar MK, et al. Photosensitivity and type I IFN responses in cutaneous lupus are driven by epidermal-derived interferon kappa. Ann Rheum Dis. 2018; 77(11): p. 1653–1664.

25. Green EK, et al. Novel ATP2A2 mutations in a large sample of individuals with Darier disease. J Dermatol. 2013; 40(4): p. 259–66.

26. Savignac M, et al. Darier disease : a disease model of impaired calcium homeostasis in the skin. Biochim Biophys Acta. 2011; 1813(5): p. 1111–7.

27. Bikle DD, et al. Calcium regulation of keratinocyte differentiation. Expert Rev Endocrinol Metab. 2012; 7(4): p. 461–472.

28. Elsholz F, et al. Calcium--a central regulator of keratinocyte differentiation in health and disease. Eur J Dermatol. 2014; 24(6): p. 650–61.

29. Matsui T, Amagai M. Dissecting the formation, structure and barrier function of the stratum corneum. Int Immunol. 2015; 27(6): p. 269–80.

30. Matsui T, et al. A unique mode of keratinocyte death requires intracellular acidification. Proc Natl Acad Sci U S A. 2021; 118(17).

31. Dickson MA, et al. Human keratinocytes that express hTERT and also bypass a p16(INK4a)-enforced mechanism that limits life span become immortal yet retain normal growth and differentiation characteristics. Mol Cell Biol. 2000; 20(4): p. 1436–47.

32. Hudson TY, et al. In vitro methods for investigating desmoplakin-intermediate filament interactions and their role in adhesive strength. Methods Cell Biol. 2004; 78: p. 757–86.

33. Huen AC, et al. Intermediate filament-membrane attachments function synergistically with actin-dependent contacts to regulate intercellular adhesive strength. J Cell Biol. 2002; 159(6): p. 1005–17.

34. Greenwald EC, et al. Genetically Encoded Fluorescent Biosensors Illuminate the Spatiotemporal Regulation of Signaling Networks. Chem Rev. 2018; 118(24): p. 11707–11794.

35. Zaver SA, et al. Live Imaging with Genetically Encoded Physiologic Sensors and Optogenetic Tools. J Invest Dermatol. 2023; 143(3): p. 353–361.E4.

36. Hiratsuka T, et al. Regulation of ERK basal and pulsatile activity control proliferation and exit from the stem cell compartment in mammalian epidermis. Proc Natl Acad Sci U S A. 2020; 117(30): p. 17796–17807.

37. Harmon RM, et al. Desmoglein-1/Erbin interaction suppresses ERK activation to support epidermal differentiation. J Clin Invest. 2013; 123(4): p. 1556–70.

38. Kam CY, et al. Desmoplakin maintains gap junctions by inhibiting Ras/MAPK and lysosomal degradation of connexin-43. J Cell Biol. 2018; 217(9): p. 3219–3235.

39. Egu DT, et al. Role of PKC and ERK Signaling in Epidermal Blistering and Desmosome Regulation in Pemphigus. Front Immunol. 2019; 10: p. 2883.

40. Khavari TA, Rinn J. Ras/Erk MAPK signaling in epidermal homeostasis and neoplasia. Cell Cycle. 2007; 6(23): p. 2928–31.

41. Simpson CL, et al. Deconstructing the skin: cytoarchitectural determinants of epidermal morphogenesis. Nat Rev Mol Cell Biol. 2011; 12(9): p. 565–80.

42. Munoz-Garcia A, et al. The importance of the lipoxygenase-hepoxilin pathway in the mammalian epidermal barrier. Biochim Biophys Acta. 2014; 1841(3): p. 401–8.

43. Crumrine D, et al. Mutations in Recessive Congenital Ichthyoses Illuminate the Origin and Functions of the Corneocyte Lipid Envelope. J Invest Dermatol. 2019; 139(4): p. 760–768.

44. Jobard F, et al. Lipoxygenase-3 (ALOXE3) and 12(R)-lipoxygenase (ALOX12B) are mutated in non-bullous congenital ichthyosiform erythroderma (NCIE) linked to chromosome 17p13.1. Hum Mol Genet. 2002; 11(1): p. 107–13.

45. Sun Q, et al. The Genomic and Phenotypic Landscape of Ichthyosis: An Analysis of 1000 Kindreds. JAMA Dermatol. 2022; 158(1): p. 16–25.

46. Russell LJ, et al. Mutations in the gene for transglutaminase 1 in autosomal recessive lamellar ichthyosis. Nat Genet. 1995; 9(3): p. 279–83.

47. Porter RM, et al. Gene targeting at the mouse cytokeratin 10 locus: severe skin fragility and changes of cytokeratin expression in the epidermis. J Cell Biol. 1996; 132(5): p. 925–36.

48. Rothnagel JA, et al. Mutations in the rod domains of keratins 1 and 10 in epidermolytic hyperkeratosis. Science. 1992; 257(5073): p. 1128–30.

49. Reichelt J, Magin TM. Hyperproliferation, induction of c-Myc and 14-3-3sigma, but no cell fragility in keratin-10-null mice. J Cell Sci. 2002; 115(Pt 13): p. 2639–50.

50. Scholl FA, et al. Mek1 alters epidermal growth and differentiation. Cancer Res. 2004; 64(17): p. 6035–40.

51. Getsios S, et al. Desmoglein 1-dependent suppression of EGFR signaling promotes epidermal differentiation and morphogenesis. J Cell Biol. 2009; 185(7): p. 1243–58.

52. Menon GK, et al. Ionic calcium reservoirs in mammalian epidermis: ultrastructural localization by ion-capture cytochemistry. J Invest Dermatol. 1985; 84(6): p. 508–12.

53. Schmidt M, et al. Ras-independent activation of the Raf/MEK/ERK pathway upon calcium-induced differentiation of keratinocytes. J Biol Chem. 2000; 275(52): p. 41011–7.

54. Carlos G, et al. Cutaneous Toxic Effects of BRAF Inhibitors Alone and in Combination With MEK Inhibitors for Metastatic Melanoma. JAMA Dermatol. 2015; 151(10): p. 1103–9.

55. See SHC, et al. Distinguishing histopathologic features of acantholytic dermatoses and the pattern of acantholytic hypergranulosis. J Cutan Pathol. 2019; 46(1): p. 6–15.

56. Fu L, et al. Targeting Extracellular Signal-Regulated Protein Kinase 1/2 (ERK1/2) in Cancer: An Update on Pharmacological Small-Molecule Inhibitors. J Med Chem. 2022; 65(20): p. 13561–13573.

57. Hovnanian A. Darier’s disease: from dyskeratosis to endoplasmic reticulum calcium ATPase deficiency. Biochem Biophys Res Commun. 2004; 322(4): p. 1237–44.

58. Huber M, et al. Abnormal keratin 1 and 10 cytoskeleton in cultured keratinocytes from epidermolytic hyperkeratosis caused by keratin 10 mutations. J Invest Dermatol. 1994; 102(5): p. 691–4.

59. Pulkkinen L, et al. Epidermolytic hyperkeratosis (bullous congenital ichthyosiform erythroderma). Genetic linkage to chromosome 12q in the region of the type II keratin gene cluster. J Clin Invest. 1993; 91(1): p. 357–61.

60. Simpson CL, Green KJ. Identification of desmogleins as disease targets. J Invest Dermatol. 2007; 127(E1): p. E15–6.

61. Hammers CM, Stanley JR. Desmoglein-1, differentiation, and disease. J Clin Invest. 2013; 123(4): p. 1419–22.

62. Dhitavat J, et al. Impaired trafficking of the desmoplakins in cultured Darier’s disease keratinocytes. J Invest Dermatol. 2003; 121(6): p. 1349–55.

63. Mauro T. Endoplasmic reticulum calcium, stress, and cell-to-cell adhesion. J Invest Dermatol. 2014; 134(7): p. 1800–1801.

64. Poulikakos PI, et al. RAF inhibitors transactivate RAF dimers and ERK signalling in cells with wild-type BRAF. Nature. 2010; 464(7287): p. 427–30.

65. Flaherty KT, et al. Combined BRAF and MEK inhibition in melanoma with BRAF V600 mutations. N Engl J Med. 2012; 367(18): p. 1694–703.

66. Awe O, et al. Drug-induced Grover’s disease: a case report and review of the literature. Int J Dermatol. 2022; 61(5): p. 591–594.

67. Anforth R, et al. Cutaneous toxicities of RAF inhibitors. Lancet Oncol. 2013; 14(1): p. e11–8.

68. Babuna Kobaner G, et al. Darier disease-like hyperkeratotic papules and invasive squamous cell carcinoma in a patient with melanoma treated with dabrafenib. Australas J Dermatol. 2018; 59(3): p. e231–e233.

69. Chu EY, et al. Diverse cutaneous side effects associated with BRAF inhibitor therapy: a clinicopathologic study. J Am Acad Dermatol. 2012; 67(6): p. 1265–72.

70. Flaherty KT, et al. Improved survival with MEK inhibition in BRAF-mutated melanoma. N Engl J Med. 2012; 367(2): p. 107–14.

71. Roskoski R, Jr. Targeting ERK1/2 protein-serine/threonine kinases in human cancers. Pharmacol Res. 2019; 142: p. 151–168.

72. David E, et al. The evolving landscape of biologic therapies for atopic dermatitis: Present and future perspective. Clin Exp Allergy. 2023; 53(2): p. 156–172.

73. Lin CP, et al. Current and emerging biologic and small molecule systemic treatment options for psoriasis and psoriatic arthritis. Curr Opin Pharmacol. 2022; 67: p. 102292.

74. Lefferdink R, et al. Secukinumab responses vary across the spectrum of congenital ichthyosis in adults. Arch Dermatol Res. 2022.

75. Kim M, et al. Transcriptomic Analysis of the Major Orphan Ichthyosis Subtypes Reveals Shared Immune and Barrier Signatures. J Invest Dermatol. 2022; 142(9): p. 2363–2374 e18.

76. Paller AS, et al. An IL-17-dominant immune profile is shared across the major orphan forms of ichthyosis. J Allergy Clin Immunol. 2017; 139(1): p. 152–165.

77. Wang H, et al. 3D cell culture models: Drug pharmacokinetics, safety assessment, and regulatory consideration. Clin Transl Sci. 2021; 14(5): p. 1659–1680.

78. Haase K, Freedman BS. Once upon a dish: engineering multicellular systems. Development. 2020; 147(9).

79. Kdimati S, et al. Patient-Derived Organoids for In Vivo Validation of In Vitro Data. Methods Mol Biol. 2023; 2589: p. 111–126.

80. Wu W, et al. Patient-derived Tumour Organoids: A Bridge between Cancer Biology and Personalised Therapy. Acta Biomater. 2022; 146: p. 23–36.

81. Novelli G, et al. Organoid factory: The recent role of the human induced pluripotent stem cells (hiPSCs) in precision medicine. Front Cell Dev Biol. 2022; 10: p. 1059579.

82. LeSavage BL, et al. Next-generation cancer organoids. Nat Mater. 2022; 21(2): p. 143–159.

83. Moore J.L. GF, Matte-Martone C., Du S., Lathrop E., Ganesan S., Shao L., Bhaskar D., Cox A., Hendry C., Rieck B., Krishnaswamy S., Greco V. G2 stem cells orchestrate time-directed, long-range coordination of calcium signaling during skin epidermal regeneration. bioRxiv. 2021: p. 10.12.464066.

84. MacLennan DH. Ca2+ signalling and muscle disease. Eur J Biochem. 2000; 267(17): p. 5291–7.

85. Britzolaki A, et al. A Role for SERCA Pumps in the Neurobiology of Neuropsychiatric and Neurodegenerative Disorders. Adv Exp Med Biol. 2020; 1131: p. 131–161.

86. Gordon-Smith K, et al. Genotype-phenotype correlations in Darier disease: A focus on the neuropsychiatric phenotype. Am J Med Genet B Neuropsychiatr Genet. 2018; 177(8): p. 717–726.

87. Bachar-Wikstrom E, Wikstrom JD. Darier Disease - A Multi-organ Condition? Acta Derm Venereol. 2021; 101(4): p. adv00430.

88. Nakajima K, et al. Brain-specific heterozygous loss-of-function of ATP2A2, endoplasmic reticulum Ca2+ pump responsible for Darier’s disease, causes behavioral abnormalities and a hyper-dopaminergic state. Hum Mol Genet. 2021; 30(18): p. 1762–1772.

89. Hua PT, Caplan JP. Darier Disease and Neuropsychiatric Illness: A Dermatologic Condition That Is More Than Skin Deep. Psychosomatics. 2020; 61(3): p. 281–283.

90. Li Pomi F, et al. Beyond the skin involvement in Darier disease: A complicated neuropsychiatric phenotype. Clin Case Rep. 2021; 9(6): p. e04263.

91. Sprecher E. What do rare and common have in common? Br J Dermatol. 2022; 187(3): p. 279–280.

92. Falchook GS, et al. Activity of the oral MEK inhibitor trametinib in patients with advanced melanoma: a phase 1 dose-escalation trial. Lancet Oncol. 2012; 13(8): p. 782–9.

93. Coulombe PA, et al. Point mutations in human keratin 14 genes of epidermolysis bullosa simplex patients: genetic and functional analyses. Cell. 1991; 66(6): p. 1301–11.

94. Cheng J, et al. The genetic basis of epidermolytic hyperkeratosis: a disorder of differentiation-specific epidermal keratin genes. Cell. 1992; 70(5): p. 811–9.

95. Mao X, et al. MAPKAP kinase 2 (MK2)-dependent and -independent models of blister formation in pemphigus vulgaris. J Invest Dermatol. 2014; 134(1): p. 68–76.

96. Berkowitz P, et al. Induction of p38MAPK and HSP27 phosphorylation in pemphigus patient skin. J Invest Dermatol. 2008; 128(3): p. 738–40.

97. Berkowitz P, et al. p38MAPK inhibition prevents disease in pemphigus vulgaris mice. Proc Natl Acad Sci U S A. 2006; 103(34): p. 12855–60.

98. Egu DT, et al. A new ex vivo human oral mucosa model reveals that p38MAPK inhibition is not effective in preventing autoantibody-induced mucosal blistering in pemphigus. Br J Dermatol. 2020; 182(4): p. 987–994.

99. Simpson CL, et al. NIX initiates mitochondrial fragmentation via DRP1 to drive epidermal differentiation. Cell Rep. 2021; 34(5): p. 108689.

100. Egolf S, et al. MLL4 mediates differentiation and tumor suppression through ferroptosis. Sci Adv. 2021; 7(50): p. eabj9141.

101. Cesinger MR, et al. The Transcriptional Regulator SpxA1 Influences the Morphology and Virulence of Listeria monocytogenes. Infect Immun. 2022; 90(10): p. e0021122.

102. Simpson CL, et al. Plakoglobin rescues adhesive defects induced by ectodomain truncation of the desmosomal cadherin desmoglein 1: implications for exfoliative toxin-mediated skin blistering. Am J Pathol. 2010; 177(6): p. 2921–37.

